# Angiogenin regulates mitochondrial stress and function via tRNA-derived fragments generation and impacting tRNA modifications

**DOI:** 10.1101/2024.02.13.580206

**Authors:** Shadi Al-Mesitef, Keita Tominaga, Abdulrahman Mousa, Thomas J Begley, Peter C Dedon, Sherif Rashad, Kuniyasu Niizuma

**Author notes:** Corresponding author. Corresponding author details: Sherif Rashad, MD, PhD.

## Abstract

Mitochondrial stress and dysfunction play an important role in many diseases, such as cancer, diabetes, and neurodegenerative diseases. We previously observed that mitochondrial electron transport chain (ETC) inhibition can induce tRNA cleavage and tsRNAs (tRNA-derived small non-coding RNAs) generation. However, whether this process is mediated via Angiogenin (ANG), the canonical enzyme responsible for tRNA cleavage, and whether it has a role in regulating the mitochondrial stress response remains to be understood. ANG is linked to Amyotrophic Lateral Sclerosis (ALS) and other conditions where mitochondrial stress plays a role in pathophysiology. Here, we aimed to examine the role of ANG in regulating the translational response to mitochondrial stress. We observed that ANG protected the cells from respiratory complex III and V inhibition specifically. Furthermore, we validated that the tsRNAs generated during mitochondrial and oxidative stress are mediated by ANG, given that their production is abrogated after ANG knock-out (KO). In addition, we observed that ANG-KO altered the tRNA modification status. Namely, we observed that ANG-KO led to the downregulation of queuosine tRNA modifications (tRNA-Q). tRNA-Q itself is related to mitochondrial translation and function. Indeed, we observed that ANG-KO led to reduced mitochondrial respiration and function. ANG altered how the cells respond to mitochondrial stress by altering the dynamic tRNA modification changes occurring during the stress response. We further examined the impact of ANG-KO on stress granules (SG) assembly as well as the knockdown of G3BP1 (core protein of SGs) on tsRNAs generation. Our results indicate that ANG regulates mitochondrial function and stress via tsRNAs generation as well as altering tRNA modifications levels. Our data also indicate that there are no direct links between tRNA cleavage and SG assembly, and both could be parallel systems for translation repression during stress.

## Introduction

Mitochondrial dysfunction is a complex phenomenon that has been associated with a wide range of abnormal conditions, including metabolic disorders, aging, cancer and neurodegenerative diseases such as Huntington’s (HD)^1^, Alzheimer’s (AD)^2^, Parkinson’s (PD)^3^, and Amyotrophic Lateral Sclerosis (ALS)^4^. Developing effective therapies requires us to understand the nuanced regulation of mitochondrial function that involves a delicate interaction of genetic, epigenetic, and environmental factors including oxidative stress that overwhelms the cells with excessive reactive oxygen species (ROS)^5^. It is becoming highly understood that mitochondrial dysfunction and stress are not singular entities. Instead, there exists a sophisticated regulatory mechanism dictating cellular responses to stress, influenced by factors like the specific mode of stress induction, including its impact on particular respiratory complexes, that play a crucial role in these responses^5^. In recent years, the tRNA epitranscriptome—a collective term used to refer to tRNA modifications, expression, and tRNA-derived small non-coding RNAs (tsRNAs)—has been recognized as a master regulator of cellular responses to stress^6–8^. Under stress conditions, site-specific nucleases cleave the mature transfer RNA (tRNA)^9^, giving rise to tsRNAs that were generally divided into tiRNAs (stress induced tRNA halves) and tRFs (tRNA derived fragments)^10,11^. The earliest reports that mention the cleavage phenomenon were in the bacterium Escherichia coli^12,13^, the unicellular eukaryote Tetrahymena^14^, the budding yeast Saccharomyces cerevisiae, and the plant Arabidopsis^15^. The discovery was extended to mammalian cells, with angiogenin (ANG) identified as the mediator of tRNA cleavage^16,17^. Under ideal growth conditions, ANG is located in both the nucleus, to promote the synthesis of the ribosomal RNA (rRNA), and the cytoplasm, where it forms a complex with the ribonuclease inhibitor 1 (RNH1). However, under diverse stressors like oxidative stress, nutrient starvation, heat shock, and viral infection, ANG becomes stimulated and mobilizes from the nucleus to the cytoplasm^18,19,17^. tsRNAs have been linked to a variety of biological roles, and it has been noted that they might be relevant to stress-related diseases and injuries. For example, in prior studies, tiRNAs were induced in vivo with renal and hepatic rodent injury models under oxidative stress^20^, in vitro after developing a hindlimb model^21^, and in an ischemia-reperfusion injury model (IRI)^22^. Moreover, 5’-tiRNAs were abundant in patients with chronic viral hepatitis^23^. Also, endogenous 5’-tiRNAs were shown to suppress translation during oxidative stress^17^. Specifically, 5’-tiRNA^Ala^ and 5’-tiRNA^Cys^ were known to mediate such inhibition by using their terminal oligonucleotide repeats which produce G-quadruplex (G4) structures that are essential for translation repression via stress granule assembly^24,25^. Previous study^10^ reported that 5’-tiRNAs under oxidative stress induced stress granule assembly (SG) in cell culture to cope with stress. Additionally, the translation silencer Y-box binding protein-1 (YB-1) did not affect translation, however, it was required for 5’-tiRNA to produce SG upon stress^26^. Furthermore, tRNA fragments (tRFs) originating from tRNA^Glu^, tRNA^Asp^, tRNA^Gly^, and tRNA^Tyr^ were found to inhibit gene expression, thereby suppressing the progression of breast cancer^27^. All these studies illustrate that tsRNAs can be considered potential biomarkers for diseases, although further exploration is needed. Despite these previous works, there have been some contradictions. For example, in cell and animal models that induced tRNA cleavage and tsRNAs generation, there was no apparent protective impact, rather, the production of tsRNAs was linked to cell death^6,22,28^. Adding to this the recent report^29^ that showed no links between tiRNAs and stress granule assembly. It is also important to highlight that the majority of studies investigating the relationship between ANG and tRNA cleavage, whether using genetic approaches to knock-out ANG or by adding recombinant ANG to cells, have primarily focused on tRNA cleavage and tsRNAs generation. However, a significant gap exists as these investigations generally overlook the direct influence of ANG on cell viability in stress response scenarios^30–32^. Additionally, the precise connections between ANG and mitochondrial function in stress, particularly in the context of the pathophysiology of diseases like ALS, where ANG is a susceptibility gene^33,34^, are yet to be explored. In this work, we aimed to elucidate the role of angiogenin-induced tsRNAs generation in regulating mitochondrial-specific stresses, via systematic inhibition of various mitochondrial respiratory complexes. We also aimed to elucidate whether there are biological links between angiogenin-induced tRNA cleavage and stress granule assembly. Our results show that ANG regulates cellular responses to specific mitochondrial stressors, that this regulation may be driven by the generation of specific subsets of tsRNAs, and that ANG is essential for mitochondrial functioning via its impact on tRNA modifications, which was not previously explored.

## Results

### Mitochondrial stress induces tsRNAs generation without stress granule assembly

We first evaluated various stressors that were known to initiate mitochondrial dysfunction and ROS production^35^. Using the wild-type HEK293T cell line, we targeted the mitochondrial complexes I, II, III, IV, and V with their inhibitors: Rotenone (RO), Thenoyltrifluoroacetone (TTFA), Antimycin A (AM), Potassium cyanide (KCN), and Oligomycin A (OLI), respectively. In addition, we used Sodium metaArsenite (AS) as a non-specific (i.e., Secondary) mitochondrial stressor. Then, we optimized the conditions of stress induction to reach 40% - 50% cell death after 4 hours of exposure to any given stressor (Fig. 1A). Next, we aimed to investigate the impact of the stressors on global protein translation, so we employed a puromycin incorporation assay. The results demonstrated a significant decrease in protein translation in the wild-type cells treated with TTFA, OLI, and AS. (Fig. 1B; Supp. 1A). Following our observation of acute translation repression in AS, we conducted a puromycin time course assay, revealing that exposure to AS leads to acute and near complete translation repression (Supp. 1B and C).Then, we evaluated the integrated stress response (ISR) and ribotoxic stress response (RSR) pathways in response to all stressors^36^. There were no significant changes when analyzing the mitogen-activated protein kinase (P38 MAPK) and its phosphorylated form (Fig. 1C and D; Supp. 1D). OLI and AS treatments led to a marked rise in the phosphorylated form of eukaryotic translation initiation factor 2a (p-EIF2a) (Fig. 1E and F; Supp. 1E). TTFA, AM, KCN, and OLI treatments exhibited a significant decrease in the phosphorylated form of the ribosomal protein S6 kinase (p-P70S6K), conversely, AS treatment resulted in a notable increase (Fig. 1G and H; Supp. 1F). Subsequently, a SYBR gold staining assay was conducted to evaluate tRNA cleavage and tsRNAs generation. The findings indicated an elevation in tsRNAs levels under AM, KCN, OLI, and AS stress compared to the control group (Fig. 1I; Supp. 1G). We examined stress granule (SG) formation upon stress by selectively targeting its core protein, GTPase-activating binding protein 1 (G3BP1)^37,38^. SGs were only observed in the AS-treated cells and barely perceptible in the RO-treated cells (less than 1% of cells). (Fig. 1J).

**Figure 1:**
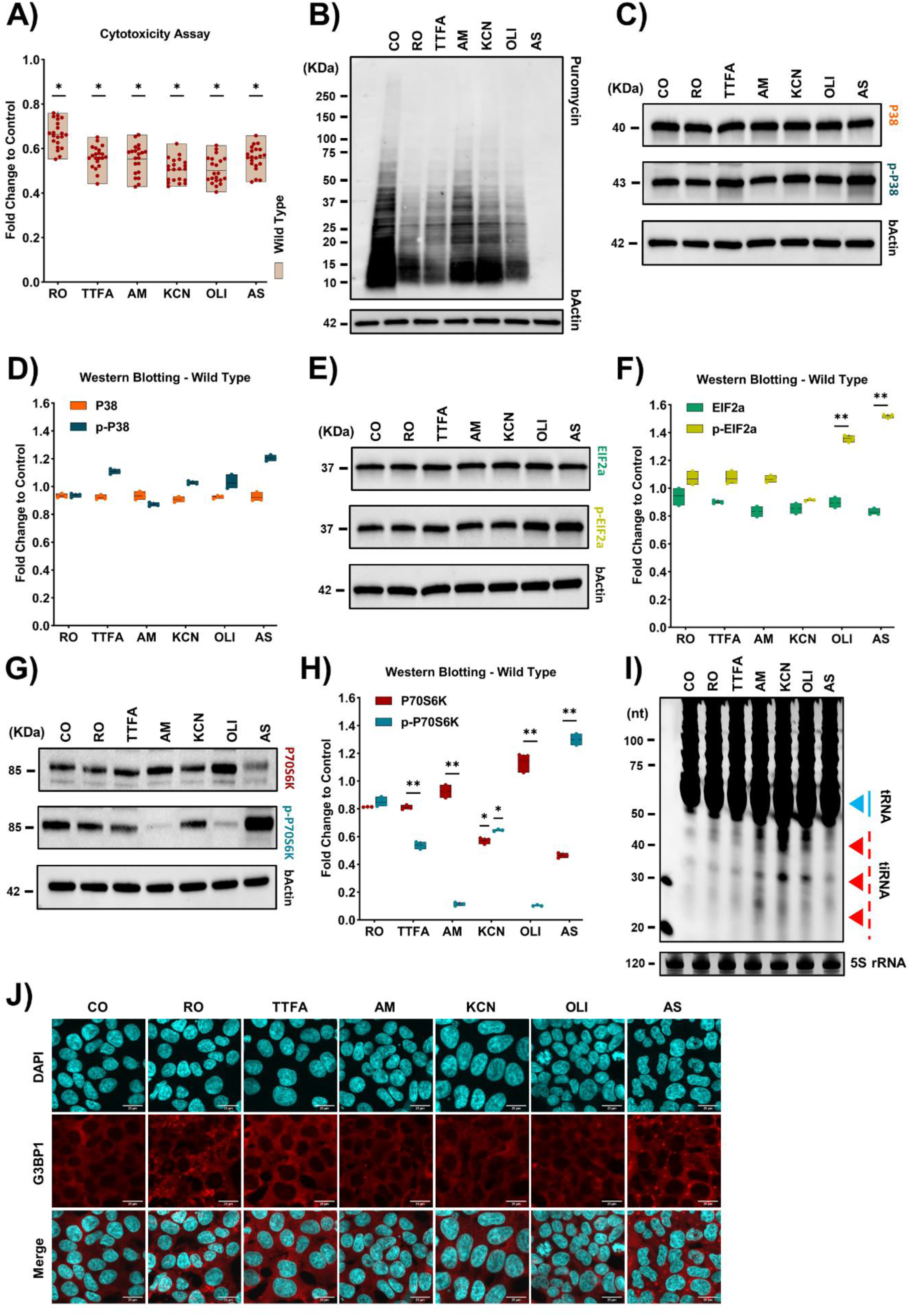
Mitochondrial stress induces tRNA cleavage. **A:** Cytotoxicity assay of various stressors. Asterisk: fold change > 1.5, *p* < 0.05. **B:** Puromycin incorporation assay after stress. **C-D:** Western blot analysis of P38 and its phosphorylated form (p-P38). **E-F:** Western blot analysis of eIF2a and its phosphorylated form (p-eIF2a). **G-H:** Western blot analysis of P70S6K and its phosphorylated form (p-P70S6K). Asterisk: fold change > 1.5, *p* < 0.05. **I:** SYBR gold staining of TBE-Urea gel electrophoresis. **J:** Immunofluorescence of G3BP1 (red), counterstained with DAPI (blue), 40X magnification, scale bar 20 μm.

### ­ Angiogenin protects against specific mitochondrial stressors

To evaluate the impact of angiogenin over-expression (ANG-OE) on cellular stress response and tsRNAs generation, we transiently transfected FLAG-tagged human angiogenin (hANG) and validated the transfection efficiency via western blotting (Fig. 2A). Then, an MTT assay revealed significant protective effects of ANG-OE against AM and OLI-induced stress on cell viability (Fig. 2B). Following that, we assessed the global protein translation. The results indicated a significant reduction in translation under the influence of AM and OLI stressors (Fig. 2C and D; Supp. 2C). Moreover, SYBR gold staining post-stress exposure revealed that ANG-OE promoted increased tRNA cleavage and tsRNAs production under TTFA, OLI, and AS stress conditions (Fig. 2E and F). Furthermore, we investigated the impact of ANG-OE on stress granule generation. Notably, SGs exhibited no significant alterations under stress conditions, mirroring the behavior observed in the wild-type cells for AS and RO (Fig. 2G).

**Figure 2:**
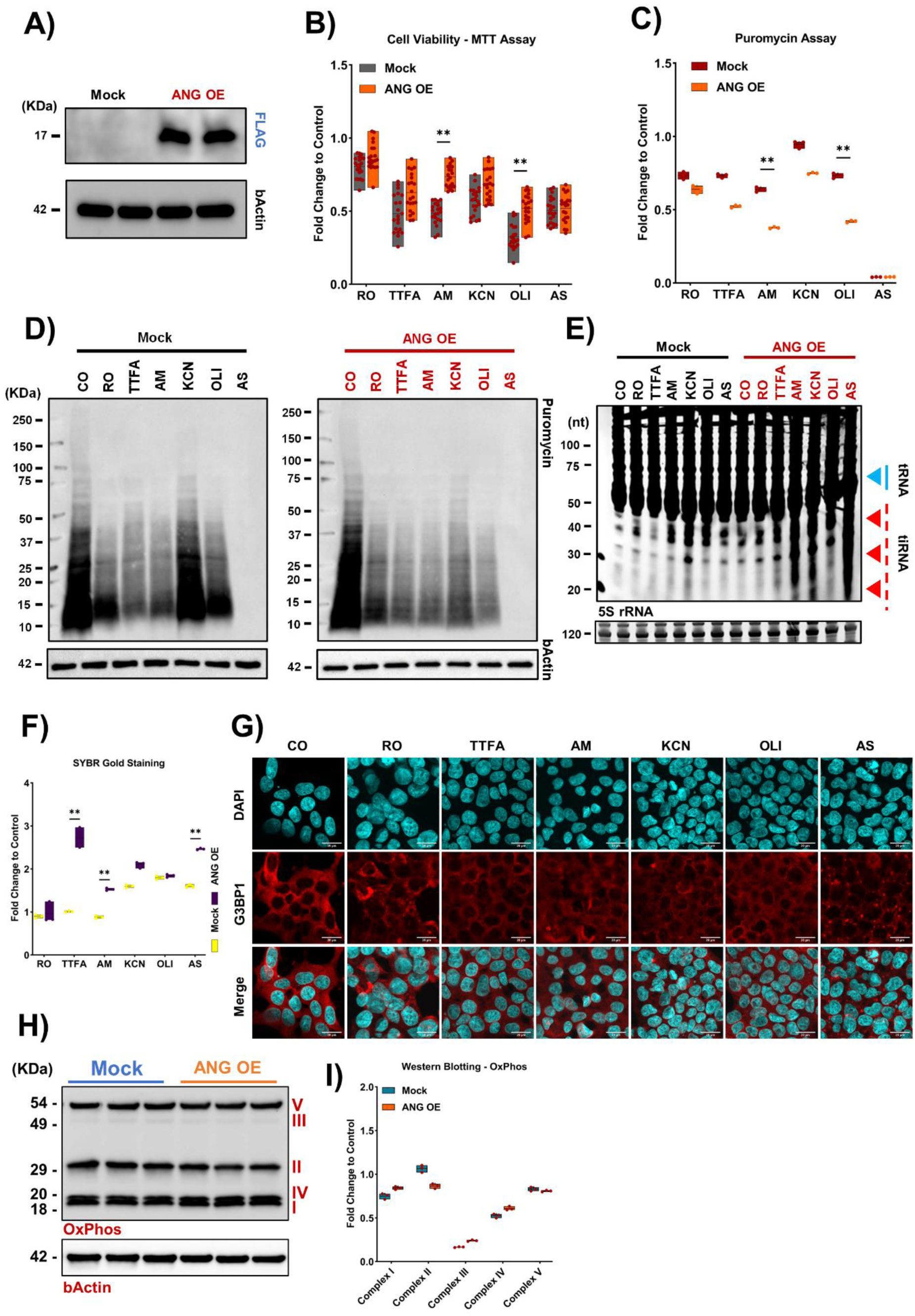
Angiogenin impacts specific mitochondrial stressors. **A:** Human angiogenin over-expression transfection efficiency via western blotting. **B:** Cell viability analysis via MTT assay after stress. Asterisk: fold change > 1.5, *p* < 0.05. **C-D:** Puromycin incorporation assay after stress. Asterisk: fold change > 1.5, *p* < 0.05. **E-F:** SYBR gold staining. Asterisk: fold change > 1.5, *p* < 0.05. **G:** Immunofluorescence of G3BP1 (red), counterstained with DAPI (blue), 40X magnification, scale bar 20 μm. **H-I:** Analysis of OXPHOS-related proteins in all mitochondrial complexes via western blotting. Asterisk: fold change > 1.5, *p* < 0.05.

For the purpose of establishing a stable angiogenin knock-out (ANG-KO) cell line, we designed a sgRNA (Supp. 2A) that targets the ANG in exonic region 2 using the LentiCRISPR-v2 vector. We confirmed the knock-out stable cell line through various techniques, including western blotting, Gene expression, and Genomic cleavage (Fig. 3A - C). Next, we evaluated the cell growth of the ANG-KO cells, revealing a reduced proliferation rate compared to Mock-KO cells (Fig. 3D). Then, we assessed the impact of ANG-KO on cell viability following stress exposure, which showed diminished viability in cells treated with AM and OLI (Fig. 3E), in agreement with the protective effect of ANG over-expression on these stresses. Afterwards, we explored the impact of ANG-KO on overall protein translation after stress exposure. All mitochondrial stressors showed a significant decrease in protein translation (Fig. 3F; Supp. 2B and C). To conduct a more in-depth analysis, we performed western blotting to observe the integrated stress response and ribotoxic stress. No significant findings after examining the P38 MAPK and its phosphorylated form (Fig. 3G; Supp. 2D - F). Exploration of the EIF2a yielded negligible findings. However, TTFA demonstrated an increase in KO cells versus the control. Additionally, OLI and AS stressors induced a marked upregulation of the p-EIF2a in both Mock and KO cells compared to the control (Fig. 3G; Supp. 2G - I). KCN and AS stressors in the KO cells induced a marked reduction in the P70S6K. Furthermore, p-P70S6K exhibited a substantial decrease with TTFA and OLI in both Mock and KO cells, AM and KCN in the KO cells, while AS exerted an opposing effect in both Mock and KO cells (Fig. 3F; Supp. 2J - L). Subsequent to SYBR gold staining, our investigation revealed a noteworthy abundance of tsRNAs generation within cells subjected to TTFA, AM, and AS treatments (Fig. 3H; Supp. 3A). Moreover, we observed no change in SG assembly status in any of the stresses after ANG-KO (Fig. 3I). To examine the mitochondrial function itself after ANG-KO, we examined the oxygen consumption rate (OCR) and the extracellular acidification rate (ECAR) through the seahorse assay. Results showed lower mitochondrial respiration in the KO cells (Fig. 3J). Notably, a significant reduction was observed in the basal and maximal respiration, proton leak, ATP production, and non-oxygen consumption rate (Supp. 3B - I). Furthermore, the oxidative phosphorylation (OXPHOS) protein expression was evaluated by performing western blotting. The results demonstrated no statistically significant changes in the KO cells compared to Mock (Fig. 3K; Supp. 3G). We also assessed the mitochondrial shapes, networks, and membrane potential in KO cells using the Mitotracker green and red immunofluorescence; however, no significant alterations were detected (Supp. 3K). Additionally, an examination of the Enhancer of mRNA Decapping 4 (EDC4), a component of processing bodies (P-bodies)^39^, revealed no observable changes (Supp. 3L).

**Figure 3:**
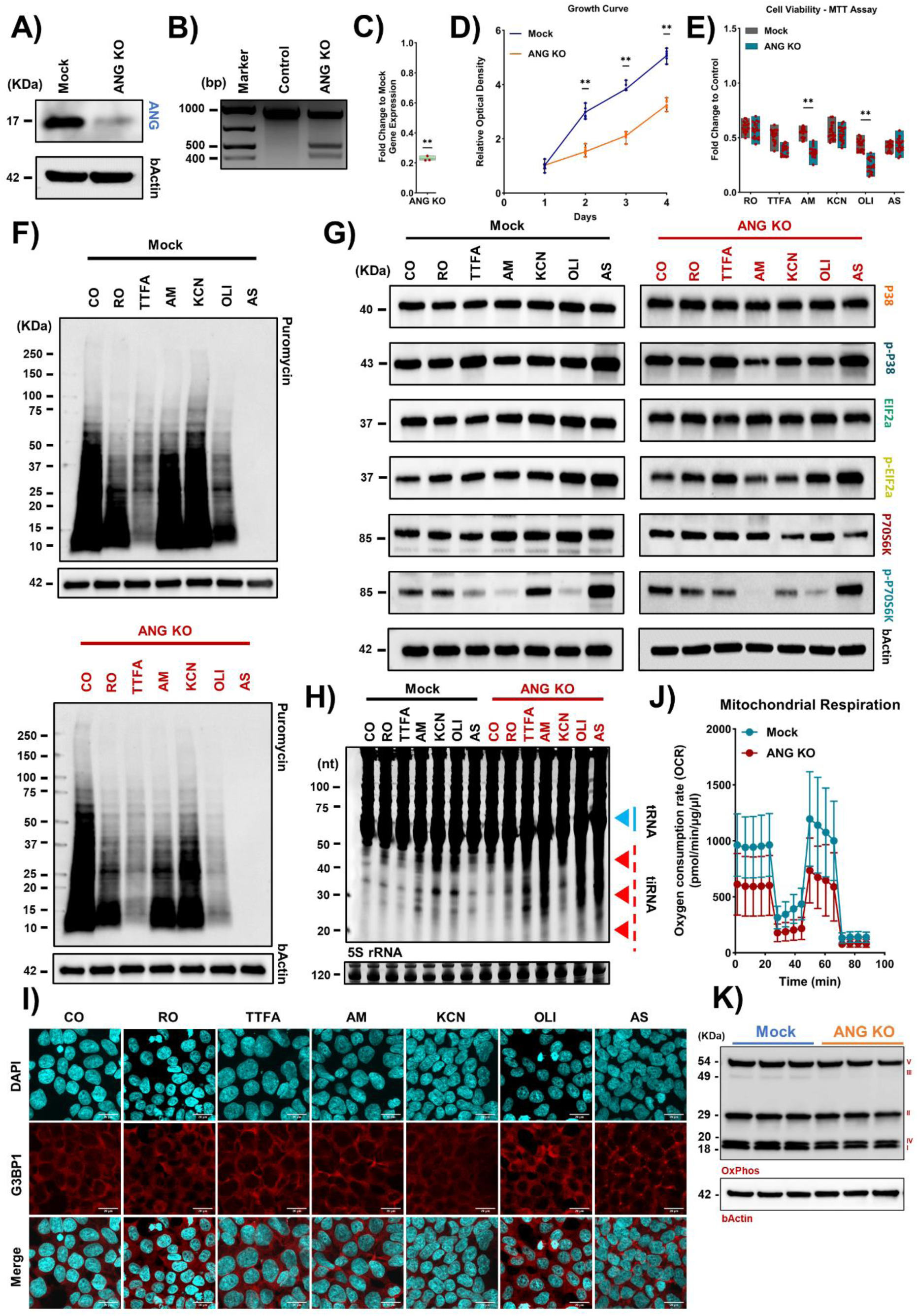
Angiogenin-KO induces translational imbalance. **A:** Validation of angiogenin KO via western blotting. **B:** Validation of ANG-KO via T7 endonuclease I assay. **C:** Validation of ANG-KO via RT-PCR. Asterisk: fold change > 1.5, p < 0.05. **D:** Cell proliferation after ANG-KO. Asterisk: fold change > 1.5, *p* < 0.05. **E:** Cell viability analysis via MTT assay after stress. Asterisk: fold change > 1.5, *p* < 0.05. **F:** Puromycin incorporation assay after stress. **G:** Analysis of ISR and RSR markers via western blotting. **H:** SYBR gold staining. **I:** Immunofluorescence of G3BP1 (red), counterstained with DAPI (blue), 40X magnification, scale bar 20 μm. **J:** Mitochondrial respiration analysis via seahorse assay. **K:** Analysis of OXPHOS-related proteins in all mitochondrial complexes via western blotting.

### ­ Angiogenin is essential for the generation of tsRNAs during stress

Given the previous data, we conducted a meticulous analysis of tsRNAs generation post-exposure to TTFA, AM, and AS stress (Fig. 4A). The small RNA sequencing results revealed no significant changes for the ANG-KO and its Mock counterpart without stress (Fig. 4B). TTFA and AM-treated cells showed major upregulation of i-tRF, 3’-tRF, and 5’-tRF fragments in Mock group, whereas ANG-KO abrogated the generation of these tRFs (Fig. 4C and D). Additionally, AS-treated cells exhibited upregulation of tiRNAs in Mock group, while ANG-KO significantly reduced tiRNAs generation after AS stress (Fig. 4E). Next, we evaluated the effect of over-expressing ANG on tsRNAs generation under the same stress conditions. Again, there was no significance when compared to its Mock under non-stress conditions (Supp. 4A). ANG over-expressing cells treated with TTFA did not show major differences in tRFs production levels (Supp. 4B). In contrast, AM-treated OE cells showed an elevation in tsRNAs generation (Supp. 4C). Meanwhile, As-treated OE cells demonstrated a downregulation of tsRNAs (Supp. 4D). Since we have two distinct systems namely OE/KO, both might have some confounding factors from the methods of generation, we aimed to isolate the actual impact of ANG on tsRNAs in response to stress via normalizing each system to its Mock then comparing them (over-expression vs Knock-out). Results revealed no difference in the absence of stress, indicating that ANG itself does not lead to spontaneous tsRNAs generation when over-expressed (Supp. 5A). However, TTFA and AM-treated cells displayed an increase in tsRNAs, namely i-tRFs and 3’-tRFs (Supp. 5B and C), while those treated with AS did not demonstrate a substantial effect, the reasons for which remain unclear (Supp. 5D).

**Figure 4:**
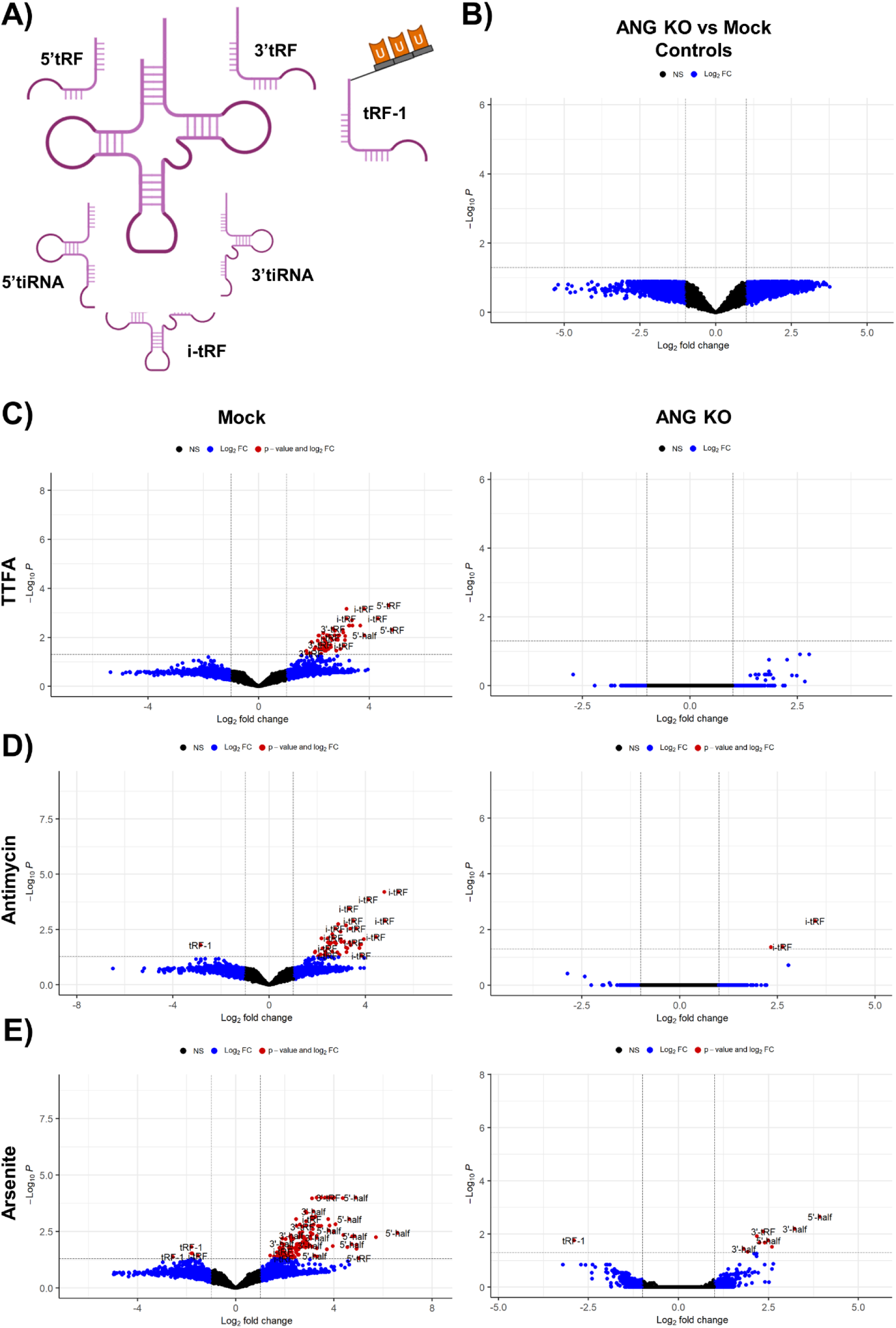
Angiogenin regulates tsRNAs generation levels. **A:** Schematic of different tsRNAs. **B-E:** Volcano plots of the Small RNA sequencing dataset for ANG-KO.

Given that tsRNAs are known to bind to RNA binding proteins (RBPs) and displace them from their target mRNAs^40^, we examined whether the tsRNAs generated during each stress might have some unique motifs that could allude to the differences observed in cell viability. We observed specific motifs that enriched in the tsRNAs generated after each stress (Supp. 6A). Given that we observed that Angiogenin was protective against Antimycin stress, and not TTFA or Arsenite, we reanalyzed the enriched motifs in antimycin vs the other stresses (Supp. 6B). This analysis revealed a set of motifs that were differentially enriched in antimycin stress. In order to further understand whether these motifs could trace to specific RBPs, we analyzed the RBPs motifs complement to the top motif in each comparison (Supp. 7A and B). In the Antimycin vs TTFA comparison, the top tsRNAs motif was complement to SNRNP70, RBM3, and MSI1. In the Antimycin vs arsenite comparison, the top tsRNAs motif was complement to RBM3, FUS, HNRNPL, and SNRNP70. This analysis revealed the potential of tsRNAs generated under different stresses to bind to specific RBPs thus altering their function. The consistency of RBM3 and SNRNP70 binding could allude to their potential contribution to the stress response, however, not enough data in the literature are present to make further assumptions.

### ­ Angiogenin regulates tRNA modifications

Given the known links between tRNA modifications and tsRNAs generation^6,32^, as well as the links between tRNA modifications and oxidative stress responses^7^, we aimed to assess the impact of tRNA modifications in response to various stressors alongside our established stable cell lines (Supp. Table 1). Initial examination of wild-type cells revealed a global downregulation of tRNA modifications in TTFA-treated cells, with notable upregulation of 5-methoxycarbonylmethyluridine (mcm^5^U) in both TTFA and AS-treated cells (Supp. 9A). Next, we explored the influence of ANG on tRNA modifications. The ANG-OE control group exhibited no changes, while the ANG-KO control group demonstrated upregulation of Wybutosine (yW) and downregulation of Queuosine (Q) and its galactosylated form (manQ) (Fig. 5A). To elucidate the causes behind such changes, we checked our RNA-seq and Ribo-seq datasets for the expression of Q and yW related enzymes, including the recently reported QTMAN and QTGAL^41,42^. QTGAL was downregulated both transcriptionally and translationally, while QTRT1 and QTRT2 did not show significant changes (Supp. 8A). We further conducted western blotting to assess the levels of QTRT1 and QTRT2 and observed significant downregulation in both enzymes at the protein levels (Supp. 8B and C). As for the yW-modifying enzymes, we did not observe a clear-cut reason for its upregulation. Further, we considered the potential that changes in mature tRNA levels could be behind the apparent changes in tRNA modifications. We revisited the small RNA-seq dataset, extracted the data on detected mature tRNAs in the ANG-KO group and the Mock-KO group, collapsed the isodecoders, and compared both groups. Our analysis revealed the downregulation of various tRNAs, most importantly, a significant downregulation of selenocysteine tRNA (tRNA-SeC^TCA^), which is known to play a role in oxidative stress response^43^, and all leucine tRNAs. However, Q or yW modified tRNAs did not exhibit significant changes (Supp. 8D). Next, we analyzed the effect of ANG over-expression on tRNA modifications changes during stress. Analysis of Mock-OE revealed upregulation of multiple modifications in the TTFA group, as well as upregulation of Q and galQ in cells treated with AM, KCN, OLI, and AS (Supp. 9B). In response to TTFA stress, ANG-OE demonstrated upregulation of mcm^5^U and Q, along with increased Q and manQ in OLI and AS-treated cells (Supp. 9C). Next, we examined the KO cells. In the Mock-KO cells, TTFA and AS stresses led to upregulation of mcm^5^U and N^6^N^6^-dimethyladenosine (m66A) respectively (Fig. 5B). ANG-KO showed a downregulation of 5-hydroxylmethylcytosine (hm^5^C) under the TTFA stress with upregulation in Q and manQ under the exposure to all stressors relative to its control (Fig. 5C).

**Figure 5:**
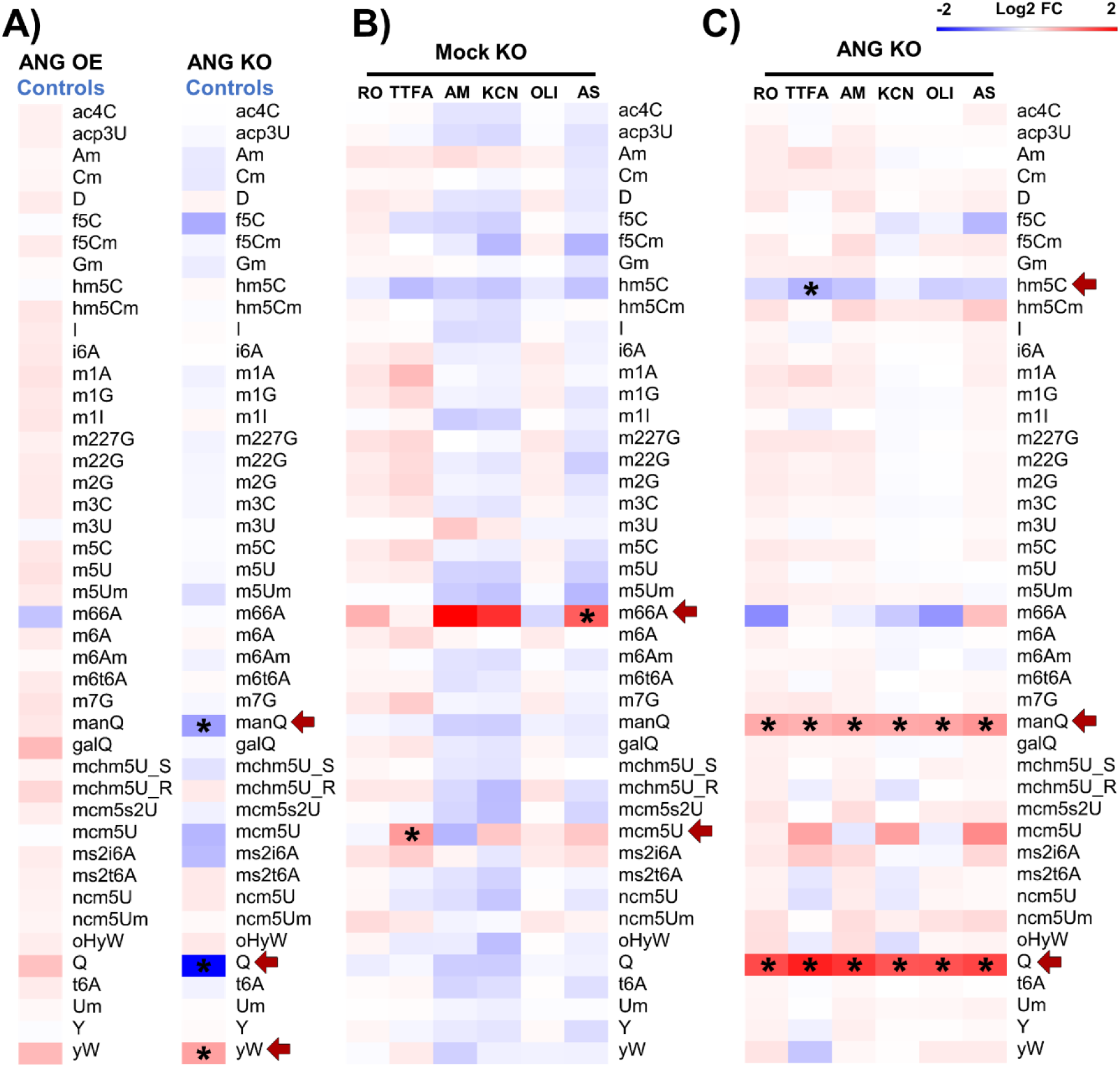
Angiogenin influences changes in tRNA modifications. **A:** Heatmaps of tRNA modifications of ANG-(OE and KO) controls. **B:** Heatmap of tRNA modifications of Mock-KO. **C:** Heatmap of tRNA modifications of ANG-KO.

### ­ Angiogenin KO alters transcription and mRNA translation

Given the impact of ANG on tRNA modifications, and its role in transcribing ribosomal RNAs, we aimed next to understand the impact of ANG-KO on mRNA levels and mRNA translation. RNA-seq Differential gene expression (DGE) analysis strong changes in the RNA levels of hundreds of genes in ANG-KO versus Mock cells (Fig. 6A). Gene ontology (GO) indicated a significant downregulation of pathways associated with mitochondrial function, such as aerobic respiration and proton motive force-driven mitochondrial ATP synthesis. There was also downregulation of translational pathways, while there was upregulation of several developmental pathways as well as protein folding related pathways (Fig. 6B; Supp. 10A). At the translational level (i.e., Ribo-seq), the protein refolding pathway was downregulated in the Ribo-seq data next to microtubule-based process, and chaperone cofactor-dependent protein refolding pathway and several chaperone and heat shock related pathways, while several stress pathways were upregulated such as endoplasmic reticulum unfolded protein response and amino acid starvation (Fig. 6C and D; Supp. 10B). We also examined the translation efficiency (TE), in which the Ribo-seq reads are normalized by the RNA-seq reads and can be an approximate proxy for translation initiation. GO pathways related to protein refolding and regulation of protein-containing complex assembly were among downregulated pathways. In contrast, endoplasmic reticulum unfolded protein response pathway and cytoplasmic translation were among the upregulated ones (Fig. 6E and F; Supp. 10C). The Pearson correlation coefficient revealed a very low correlation between RNA-seq and Ribo-seq datasets (*R* = 0.14) and a strong inverse correlation between translation efficiency and RNA levels (*R* = -0.67) suggesting the presence of strong translational dysregulation after ANG-KO (Fig. 6G).

**Figure 6:**
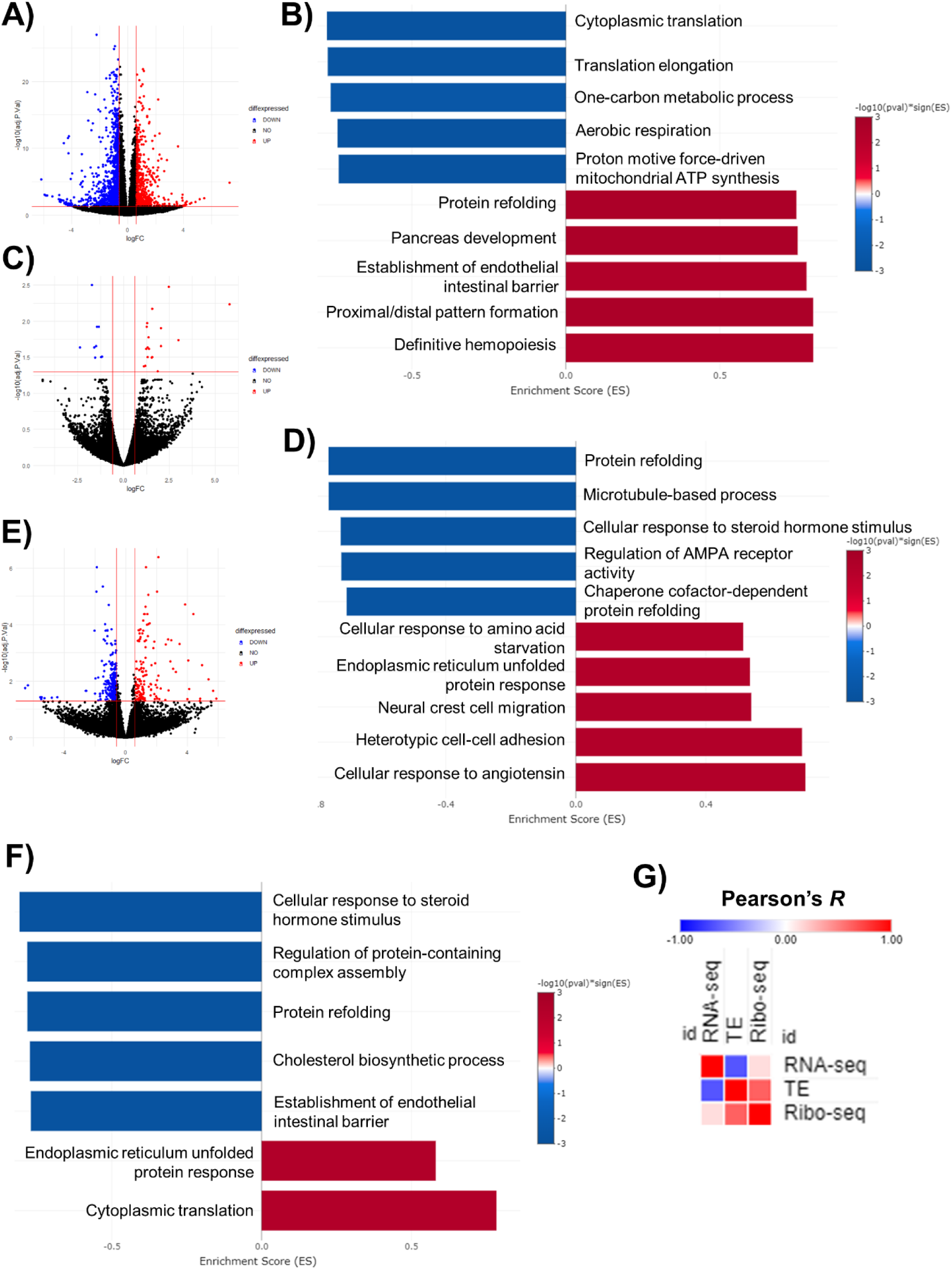
Angiogenin-KO upregulates pathways associated with mitochondrial dysfunction. **A:** Volcano plot of the RNA sequencing dataset. **B:** GOBP analysis of the RNA sequencing dataset. **C:** Volcano plot of the Ribosome profiling dataset. **D:** GOBP analysis of the Ribosome profiling dataset. **E:** Volcano plot of the Translation efficiency dataset. **F:** GOBP analysis of the Translation efficiency dataset. **G:** Pearson’s correlation analysis between different datasets.

In order to further understand the translational impact of ANG-KO on the cells, we examined the ribosome occupancy across the codon sequence in our Ribo-seq data (Fig. 7A - C). Across the coding sequence, there were changes in ribosome occupancy compared to Mock cells. In addition, most ribosome protected fragments (RPF) reads were stacked towards the start codon in ANG-KO cells compared to the end codon, indicating potential translational dysregulation. Notwithstanding, we did not observed changes in the A-site ribosome dwelling across various codons, despite the previously observed changes in tRNA modifications (Fig 7D). Next, we analyzed the isoacceptors codon frequencies and total codon frequencies in the translated mRNAs^44,45^ using the Ribo-seq dataset as well as translational efficiency data. There was a clear codon biased translation in the Ribo-seq dataset that was evident in both analyses. Importantly, we observed underusage of His^CAC^ codon compared to its synonymous His^CAT^, both of which are decoded by Queuosine modifications. However, other Q-codons (Tyr, Asp, and Asn) showed a clear preference for the NAC codons compared to NAT codons (Fig 7E and F). At the level of translation efficiency, we did not observe a strong codon biased change, indicating that the changes in TE are mostly not driven by tRNA modifications changes (Supp. 11A and B).

**Figure 7:**
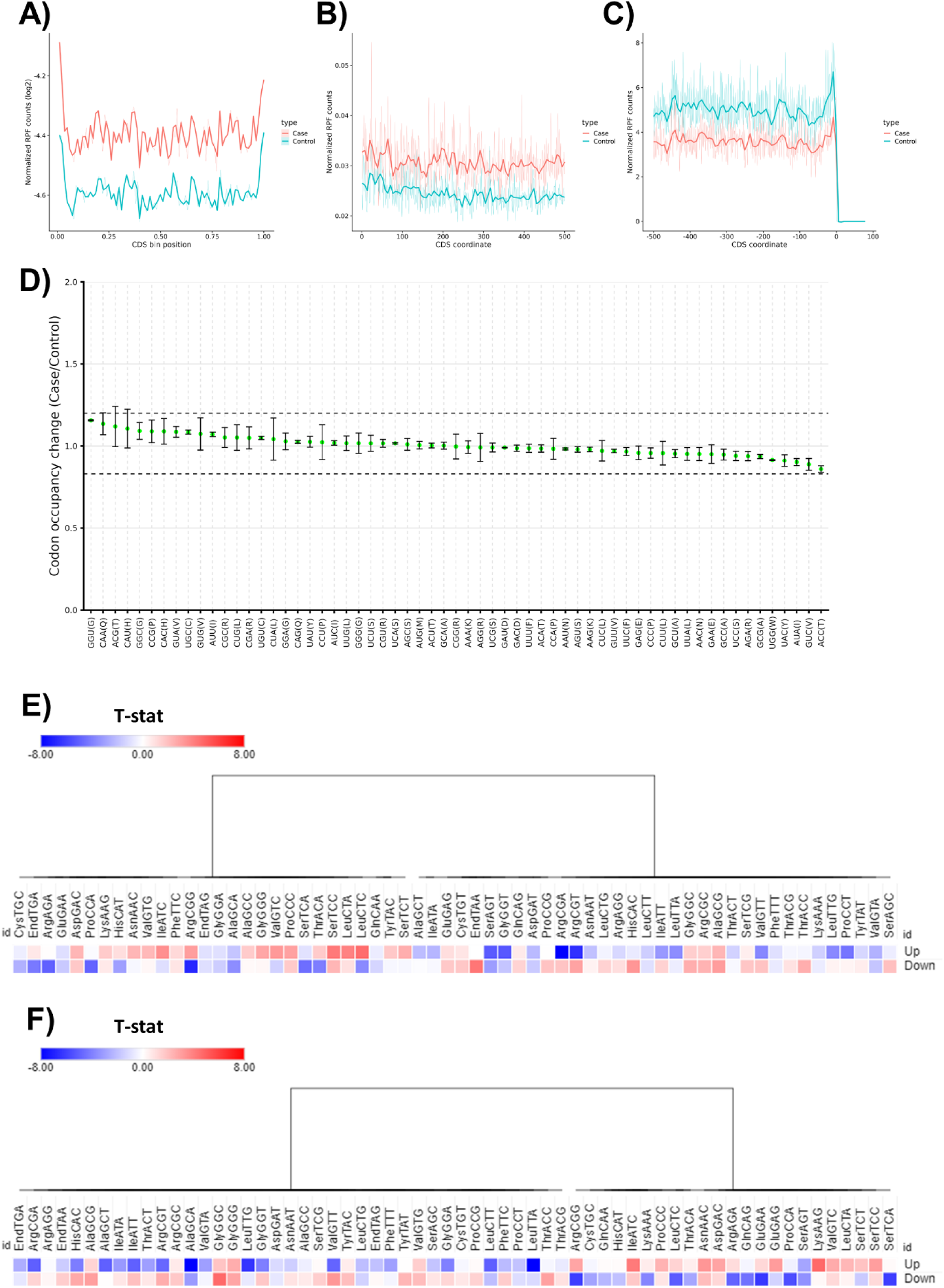
Angiogenin-KO affects mRNA translation. **A-C:** Occupancy metagene plots in ANG-KO cells (Across CDS, downstream from start codon, and upstream from end codon respectively). **D:** Ribosome A-site pausing in ANG-KO. **E:** Isoacceptors codon frequency analysis of the Ribosome profiling dataset. **F:** Total codon count analysis of the Ribosome profiling dataset.

### ­ Targeting stress granules accentuates stress-induced tsRNA production

In pursuit of understanding the links between tsRNAs and stress granules, two short hairpin RNAs (shRNAs) were designed (Supp. 12A and B) to selectively target GTPase-activating binding protein 1 (G3BP1), a vital stress granule component^37,38^. The efficacy of the knock-down (KD) was verified through western blotting (Fig. 8A). Subsequent evaluation of cellular proliferation revealed attenuated growth in KD cells compared to Mock (Fig. 8B). Despite exposing cells to diverse stressors, no significant differences in cell viability emerged (Fig. 8C). Notably, a puromycin incorporation assay demonstrated significantly reduced global translation rates in TTFA and KCN-treated KD cells (Fig. 8D and E). Additionally, utilizing SYBR gold staining, we investigated the impact of G3BP1-KD on tsRNAs production post-stress, revealing an unexpectedly heightened effect on tRNA cleavage and tsRNAs generation across all stress conditions and significant in OLI and AS-treated KD cells (Fig. 8F and G).

**Figure 8:**
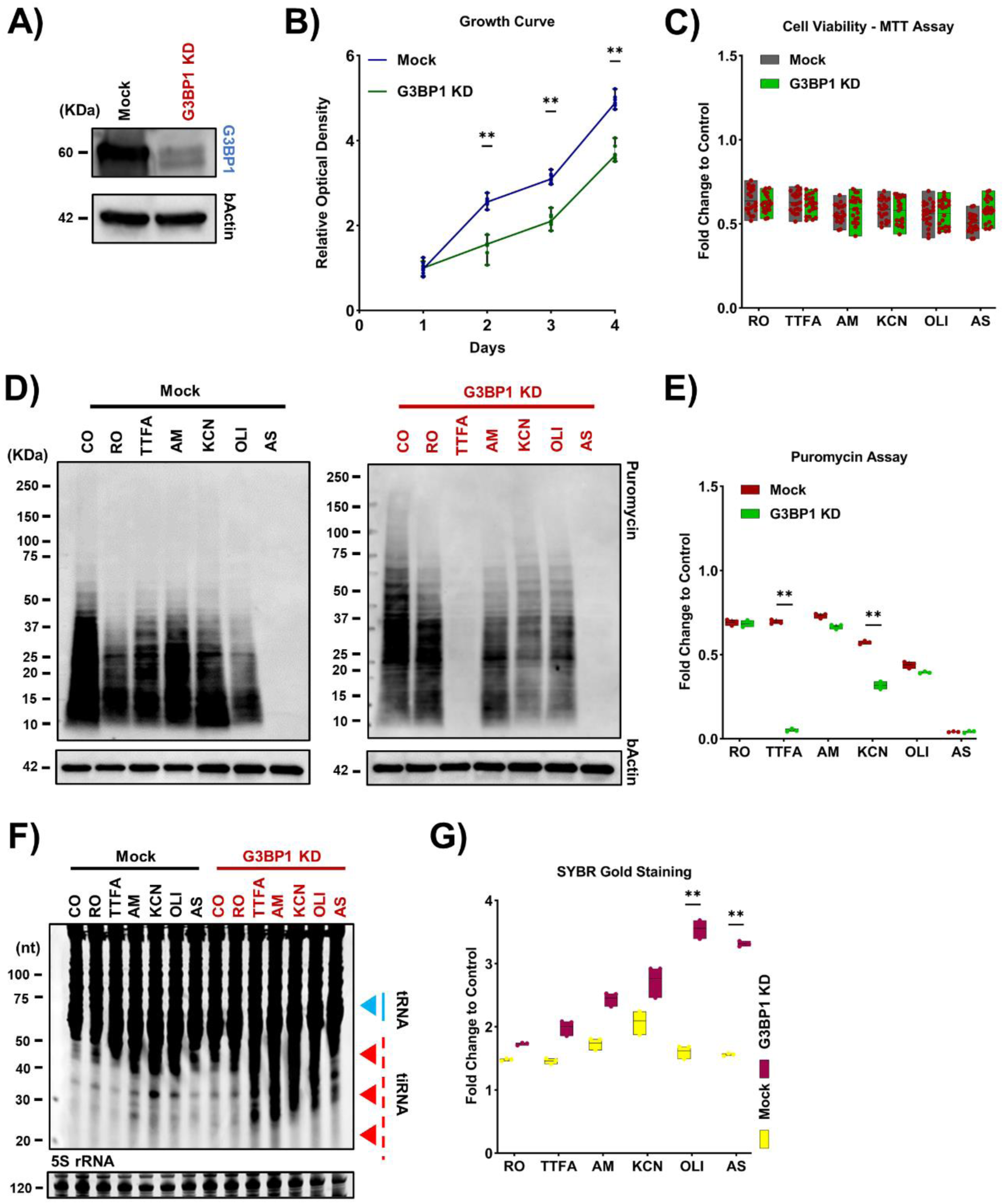
G3BP1-KD upregulates tsRNAs generation. **A:** Validation of G3BP1-KD via western blotting. **B:** Cell proliferation after G3BP1-KD. Asterisk: fold change > 1.5, *p* < 0.05. **C:** Cell viability analysis via MTT assay after stress. **D-E:** Puromycin incorporation assay after stress. Asterisk: fold change > 1.5, *p* < 0.05. **F-G:** SYBR gold staining. Asterisk: fold change > 1.5, *p* < 0.05.

## Discussion

In this study, we show that tsRNAs are generated during mitochondrial stress as well as general oxidative stress. We also show that ANG plays a critical role in generating these tsRNAs; however, the mechanisms of tsRNAs production are stress dependent. We also show that ANG plays a role in protecting the cells against specific mitochondrial stresses, such as respiratory complex III and V inhibition. We further employed the largest tsRNAs sequencing experiment of its kind, to the best of our knowledge, to probe the dynamics of tsRNAs generation as governed by ANG. Our analysis revealed that ANG-KO, or over-expression in itself, does not induce tsRNAs; however, ANG did impact tsRNAs during stress. Further, we show that the tsRNAs generated during different stresses encode different motifs that can bind to different RBPs, thus impacting their function. While it is too early to make further assumptions about this process, such data provide exciting future directions to explore the diverse and specific tsRNAs activities. We further show, for the first time, that ANG can influence tRNA modifications levels. In particular, we show that ANG-KO reduced Queuosine (tRNA-Q) and mannosyl-queuosine (manQ) levels. Given the links between tRNA-Q and mitochondrial function^41,42^, it can explain the ANG-KO-induced mitochondrial dysfunction observed herein. Interestingly, such changes occurred without alterations in the levels of tRNA-Q-modified mature tRNAs, while the tRNA-guanine transglycosylase (TGT) complex protein levels were reduced. We also show that ANG-KO induced various transcriptional and translational changes and induced translational stress in the cells. Finally, our data revealed no direct relation between tsRNAs generation and SG assembly, in agreement with the recently published work^29^. However, our data points to the possibility that SG assembly and tsRNAs generation are two parallel mechanisms that the cell can use to induce translation repression in response to stress. Angiogenin, a 123-amino acid single-chain protein initially isolated from the adenocarcinoma cell line (HT-29)^46^, belongs to the RNase-A superfamily^47^ and is recognized for its role in promoting angiogenesis^48^. Beyond its angiogenic function, ANG has been associated with neurodegenerative diseases^49^, particularly amyotrophic lateral sclerosis (ALS). Early studies identified a mutation in sporadic ALS patients, suggesting a potential decrease in ANG functions^50^. Furthermore, loss-of-function mutations in the angiogenin gene have been linked to familial cases of ALS and Parkinson’s disease (PD)^33^. Additionally, delivering human ANG to mice exhibiting ALS-like symptoms has demonstrated an extension of lifespan and improved motor function^51^. Moreover, elevated levels of the 5’ValCAC subset of tRNAs, cleaved by angiogenin, have been observed in both ALS mouse models and patients^52^. Increased 5’ValCAC levels correlate with an enhanced angiogenin-mediated stress response, contributing to a decelerated disease progression in ALS. It is noteworthy to mention that the majority of these studies employed ANG recombinant protein for the examination of tRNA cleavage. On top of that, ANG was found significantly upregulated in human malignant melanoma and cervical cancer cell lines under stress conditions^53,54^, and associated with metastasis in colorectal cancer (CRC) patients^48^. Recent findings^29^ have tested the claims regarding tsRNAs inducing stress granule assembly and providing protection against oxidative stress caused by AS stress, as previously reported by^10,25^. The study examined the impact of various stressors (AS, H2O2, nutrient starvation) on wild immortalized and primary cell lines. Contrary to prior beliefs, their findings align with ours, indicating no correlation between tsRNAs and stress granules assembly. Notably, stress granules were observed with lower AS doses that minimally induce tsRNAs generation. In addition, they replicated^29^ an experiment from^10^ by transfecting synthetic 5’-tiRNAs into cells and assessing their effect on stress granule assembly post-AS stress exposure. Surprisingly, even at supra-physiologic copy numbers, 5’-tiRNAs failed to induce stress granule formation. These results, coupled with our own, raise doubts about the direct role of 5’-tiRNAs in inducing stress granule assembly during the stress response. Furthermore, protecting tRNAs from cleavage by altering their 1-methyladenosine (m1A) status was effective in guarding against Arsenite-induced oxidative stress^6^. Collectively, it appears that tRNA cleavage might be a last-ditch attempt of the dying cells to induce translation repression to respond to an overwhelming stress. It also appears that SG assembly and tsRNAs generation are not directly connected, but two parallel processes that the cells can tap into to regulate translation during stress. However, more work is needed to fully comprehend such connections. In this work, we present, for the first time, a high throughput analysis of tRNA modifications in response to various mitochondrial stresses. While previous works have shown the dynamic nature of tRNA modifications in response to oxidative stress, such works were limited to general oxidative stresses such as Arsenite or H2O2^7,55^. The links between mitochondrial stress and tRNA modifications, and how such links influence the translational responses to mitochondrial stress were unexamined. This is particularly important given that the mitochondria do not possess stress response genes, rather, the mitochondrial stress has to be relayed to the cytosol^56^. In the wild-type cells, we observed stress specific changes in tRNA modifications levels. Interestingly, TTFA stood out as it induced the downregulation of a large number of tRNA modifications. We also observed an important technical aspect of such analysis. Namely, gene over-expression or KO both altered how cells responded to stresses at the tRNA levels compared to wild-type cells. This could be the result of how the cells are manipulated to generate gene editing, altimately impacting the cellular translatome and epitranscriptome. Thus, such changes and differences should be kept in mind when interpreting data from transgenic cells. Our work also shows for the first time the links between ANG and tRNA modifications. Given the links between tRNA modifications and many diseases such as cancer^57^, diabetes^58^, and in particular neurodegenerative diseases such as ALS^59^, such connection between ANG would have great implications in diseases in which it plays a role. ANG-KO impacted queuosine, which is known to impact mitochondrial function as well as response to oxidative stress^7,42^. Q also links ANG-KO to the tetrahydrobiopterin pathway (BH4 pathway) which plays an important role in neurodegenerative diseases and its dysregulation can impact mitochondrial function^60–62^. This connection provides a further link between ANG and mitochondrial function as well as an interesting link to neurodegeneration via metabolic dysregulation of the BH4 pathway. Indeed, BH4 pathway was shown to be involved in ALS pathophysiology^63^. In addition, ANG-KO or over-expression altered the cellular tRNA modification changes in response to mitochondrial stresses. In summary, our work shows that ANG impacts mitochondrial function and stress response via a complex interaction of tsRNAs and tRNA modifications. Such complexity was not previously revealed, and while it makes for an interesting case in ANG functionality, it will make the isolation of specific downstream effects of ANG more challenging. We also show that tsRNAs generation is not a haphazard process, and that the motifs enriched in tsRNAs during stress are highly diverse. It remains to be seen what the regulators and rules are governing such specificity as it was not previously explored. While our work shows interesting new mechanisms of how ANG works, there remains a gap in our understanding of the translational role of ANG, and more analysis is needed. In particular, what other factors influence tsRNAs generation in addition to ANG, such as specific tRNA modifications, or other unknown factors need to be studied. Also, this analysis was conducted in HEK293T cells. While the analysis offers interesting clues to the link between ANG and pathologies such as ALS, HEK293T cells are not representative of such diseases, and more analysis needs to be performed in a disease-relevant cells, such as neurons in case of neurodegenerative diseases, to achieve a better understanding of the pathologies at hand.

## Materials and methods

### Cell culture

The human embryonic kidney (HEK) 293T cell line was purchased from Takara (Cat# 632180) and cultured in a humidified 95% air / 5% CO2 incubator at 37°C in a standard Dulbecco’s modified Eagle’s medium (DMEM) (Gibco™, Cat# 10566-016) supplemented with 4.5 g/L D-Glucose, 10% Fetal bovine serum (FBS) (Corning, Cat# 35-079-CV), and 1% Sodium pyruvate (Nacalai Tesque, Cat# 06977-34). No antibiotics were added to the growth media. For passaging, cells were treated with 0.25% Trypsin-EDTA (Gibco™, Cat# 25200-072). As for counting, Trypan blue stain 0.4% (Gibco™, Cat# 15250-061) and Disposable hemocytometer (Funakoshi, Cat# 521-10) were used.

### Cell stress experiment

A density of 5 × 10^4^ cells/well was seeded in a 96-well plate (Corning Incorporated, Cat# 3596), as well as 1 × 10^6^ cells/well in a 6-well plate (Falcon, Cat# 353046) respectively, one day in advance to reach 80-90% confluency at the time of the experiment. Then, a fresh medium containing the indicated six concentrations down below was added to the cells for a duration of four hours.

1. Exposure to Rotenone (RO) (100 mM stock) (Sigma Aldrich, Cat# R8875-1G), mitochondrial complex I (NADH dehydrogenase) inhibitor. Fresh medium containing a concentration of 80 µM was added to the cells.
2. Exposure to Thenoyltrifluoroacetone (TTFA) (100 mM stock) (Abcam, Cat# ab223880), mitochondrial complex II (Succinate dehydrogenase) inhibitor. Fresh medium containing a concentration of 1.5 mM was added to the cells.
3. Exposure to Streptococcal antimycin A (AM) (2 mg/mL stock) (Sigma Aldrich, Cat# A8674-50MG), mitochondrial complex III (Cytochrome c reductase) inhibitor. Fresh medium containing a concentration of 50 µg/mL was added to the cells.
4. Exposure to Potassium cyanide (KCN) (1 M stock) (Sigma Aldrich, Cat# 60178-25G), mitochondrial complex IV (Cytochrome c oxidase) inhibitor. Fresh medium containing a concentration of 15 mM was added to the cells.
5. Exposure to Oligomycin A (OLI) (1 mg/mL stock) (Abcam, Cat# ab141829), mitochondrial complex V (ATP synthase) inhibitor. Fresh medium containing a concentration of 20 µM was added to the cells.
6. Exposure to Sodium meta-Arsenite (AS) (1 M stock) (Sigma Aldrich, Cat# S7400-100G), non-mitochondrial and endoplasmic reticulum stressor. Fresh medium containing a concentration of 600 µM was added to the cells.

### Cell viability assay

For measurement of the cell viability, MTT colorimetric assay (Fujifilm, Cat# SS-8190) was conducted in a 96-well plate following the stress experiment.

### Cell proliferation assay

Cells were seeded at a low density in a 96-well plate (10,000 cells per well) and cultured for a week, while cell proliferation was evaluated and measured using the MTT assay.

### Over-expression of human angiogenin (ANG-OE)

A density of 1 × 10^6^ cells/well was cultured for 24 hours in a 6-well plate. Then, FLAG-tagged human angiogenin plasmid (hANG) (OriGene, Cat# RC208874) and Mock plasmid were transiently transfected according to the manufacturer’s protocol into HEK293T cells in full growth medium using Lipofectamine 3000 (Thermo Fisher Scientific, Cat# L3000-015). To assess the transfection efficiency, western blotting was performed three days after the transfection.

### Angiogenin knock-out (ANG-KO)

Lentivirus CRISPR oligonucleotides targeting the exonic regions of the angiogenin gene (F: GATACTGTGAAAGCATCATG) and a non-targeting control (F: GTTCCGCGTTACATAACTTA) were designed using the BROAD Institute CRISPick design tool. They were cloned into lentiCRISPR-v2 (Addgene, Cat# 52961) following the GeCKO protocol with slight modifications^64^. Briefly, the lentiCRISPR-v2 vector was digested by BsmBI-v2 (NEB, Cat# R0739L) and dephosphorylated by Quick CIP (NEB, Cat# M0525S). Then, the synthesized phosphorylated oligos were annealed in a thermocycler (Thermo Fisher Scientific). Next, the annealed oligos were ligated into the digested lentiCRISPRv2 backbone using the DNA ligation kit (Mighty Mix) (Takara, Cat# 6023). Afterward, the ligated DNA was transformed into One Shot Stabl3 competent cells (Thermo Fisher Scientific, Cat# C7373-03) and selected on LB agar plates with 100 µg/mL Ampicillin sodium (Fujifilm, Cat# 012-23303), followed by verifying the correct incorporation of the sgRNA target sequence into the lentiCRISPR-v2 vector. Following this procedure, the HEK293T cells were transfected with the respective constructs along with envelope plasmid (pMD2.G) (Addgene, Cat# 12259), and packaging plasmid (psPAX2) (Addgene, 12260) using Lipofectamine 3000 according to the manufacturer’s protocol. Thereafter, the 293T media containing lentivirus was collected, filtered through a 0.45 µm Millex-HV filter (Merck, Cat# SLHV033RS), and infected the 293T cells with 8 µg/ml polybrene (Sigma Aldrich, Cat# TR-1003-G) for 24 hours. By using 2 µg/mL of Puromycin (Sigma Aldrich, Cat# P8833) over a week, stably infected cells were harvested and isolated through limiting dilution. Finally, single cells were expanded, and the level of the knock-out was evaluated by western blotting, T7 endonuclease I assay kit following the manual’s protocol (GeneCopoeia, Cat# IC005) (F: ATGGGGAAGAACGGTTGGAG), (R: GGGGGAAAGATCAATATGCCCA), and gene expression by RT-PCR.

### shRNA knock-down of G3BP1

A set of two human G3BP1-specific shRNAs (#1 F: GATGCTCATGCCACGCTAAAT) (#2 F: AGTGCGAGAACAACGAATAAA) and a non-targeting control (F: GCACTACCAGAGCTAACTCAGATAGTACT) were designed using the BROAD Institute design tool and cloned into pLKO.1 vector (Addgene, Cat# 8453), followed by validating the successful insertions using Colony-PCR and Sanger sequencing. Next, pLKO.1-inserted shRNAs were co-transfected with envelope plasmid (pMD2.G), and packaging plasmid (psPAX2) by using Lipofectamine 3000 according to the manufacturer’s protocol in HEK293T cells. Then, the 293T cells were infected with lentivirus in the presence of 8 µg/ml polybrene for 48 hours. Puromycin selection was performed over a week by adding fresh medium containing 2 µg/mL of Puromycin. Finally, puromycin-resistant cells were selected and the level of G3BP1 was examined by western blotting.

### RNA isolation and Quality control

Following the stress experiment, cells were lysed with QIAzol lysis reagent (QIAGEN, Cat# 79306), and Total RNA was isolated using miRNeasy mini kit (QIAGEN, Cat# 217004) according to the manufacturer’s protocol and as previously described^22,65^. Small RNA was extracted using Purelink miRNA isolation kit (Thermo Fisher Scientific, Cat# K157001) following the manufacturer’s instructions. Assessment of RNA purity and concentration was examined using Nanodrop (Thermo Fisher Scientific, Cat# ND-ONE-W), while the RNA integrity number (RIN) was determined via the RNA 6000 Nano Kit (Agilent, Cat# 5067-1511) on the Agilent Bioanalyzer 2100. Samples with RIN of ≥ 9 were used.

### SYBR gold staining

SYBR gold staining was performed as previously mentioned in^6^. Briefly, 3 µg of total RNA were mixed with Novex TBE-Urea sample buffer (Thermo Fisher Scientific, Cat# LC6876) and heated at 70°C for 3 minutes. Following that, samples were loaded into Novex 10% TBE-Urea gels (Invitrogen, Cat# EC68755BOX) in 1X TBE buffer (Bio-Rad, Cat# 1610733). Next, gels were then incubated with 10,000-fold diluted SYBR Gold nucleic acid gel (Life Technologies, Cat# S11494) in 1X TBE buffer for 30 minutes at room temperature (RT). Finally, stained gels were visualized using a ChemiDoc machine (Bio-Rad).

### RNA sequencing (RNA-seq)

Two biological replicates per group were used for RNA-seq library preparation, utilizing the NEBNext Poly (A) mRNA Magnetic Isolation Module (NEB, Cat# E7490) to enrich mRNA and the Ultra II Directional RNA Library Prep Kit (NEB, Cat# E7760) in accordance with the manufacturer’s instructions. Library quality was evaluated using the Agilent DNA 1000 kit (Agilent, Cat# 5067-1504) on the Agilent Bioanalyzer 2100, while library concentration was determined using the NEBNext Library Quant Kit for Illumina (NEB, Cat# E7630). All input RNA had RNA integrity number (RIN) of > 7. Following pooling, the libraries underwent sequencing by Macrogen on the Illumina HiSeq X-ten platform (150bp paired end).

### RNA-seq data analysis

Quality control for Raw Fastq was performed using FastQC. Raw reads were then trimmed with Trimmomatic^66^ and adaptor sequences and low-quality read removed. Reads were aligned to the human reference genome hg38 (downloaded from UCSC genome browser) using the splice aware aligner HISAT2^67^. Mapped reads BAM file were then counted to gene features by FeatureCounts^68^ with standard settings. Differentially expressed genes (DEGs) were analyzed using Limma-Voom^69^ with normalization method TMM.

### Small RNA sequencing

The Arm-seq method, as detailed^70^, was employed with few adjustments. In summary, 500 ng of RNA underwent deacylation in a 0.1 M pH 9.0 Tris-HCL buffer at 37°C for 1 hour to eliminate 3’ conjugated amino acids. Subsequently, the deacylated RNA was subjected to AlkB demethylase treatment (rtStar tRNA-optimized First-Strand cDNA Synthesis Kit, ArrayStar, Cat# AS-FS-004) at RT for 2 hours to mitigate tRNA methylation. Purification of RNA employed the Zymo Directzol RNA micro kit (Zymo Research, Cat# R2060), followed by end repair with T4 Polynucleotide Kinase (T4 PNK; NEB, Cat# M0201) at 37°C for 30 minutes. Library preparation utilized the NEBNext Multiplex Small RNA Library Prep Set for Illumina (NEB Cat# E7300/7580) according to the manufacturer’s protocol. Size selection of the amplified cDNA library occurred through 6% TBE PAGE within a range of 140 to 210 bp (microRNA to mature tRNA). Assessment of library concentration and quality was conducted using the Bioanalyzer 2100 with a DNA 1000 kit. Sequencing was performed on the Illumina Hiseq-X ten instrument in a pair-end, 150 bp read format, pooling two biological replicates per group.

### Small RNA-seq data analysis

Small RNA-seq data analysis was conducted as previously reported^44^. In short, Seqprep was used to remove adapters and collapse paired end-reads into single files. Mintmap V2 was used to analyze tsRNAs^71^. tRAX was used to analyze mature tRNA expression^31^. Motif analysis was conducted on the top enriched tsRNAs in the Mock-KO group (FDR < 0.05, Log2FC > 1) using MEME-suite^72^ with the parameters: ZOOPS, minimum width = 6, maximum width = 8, minimum sites = 2, and maximum number of motifs = 5.

### Ribosome profiling (Ribo-seq)

Ribosome profiling was conducted in accordance with our prior work^73^, adapting a protocol with minor adjustments as outlined by^74^. Subsequent to sample collection, ribosome foot-printing involved the addition of 1.25 U/μL RNase I (NEB Cat# M0243L) to 500 μL clarified lysate, followed by 45 minutes incubation at RT on a rotator mixer. TRIzol reagent facilitated RNA extraction using the Qiagen miRNeasy kit. Ribosome protected fragments (RPFs) within the range of 27-35 nucleotides (nt) were selected through TBE-Urea gel electrophoresis. rRNA depletion was achieved using the NEBNext rRNA depletion kit v2 (NEB, Cat# E7400L), and subsequent end-repair, post-purification of rRNA-depleted samples, was performed using the Zymo Oligo Clean and Concentrator Kit (Zymo Research, Cat# D4060). Sequencing libraries for ribosome profiling were prepared following the NEBNext Multiplex Small RNA Library Prep kit for Illumina’s protocol (NEB, Cat# E7300S). Ribo-seq, generating pair-end sequencing reads of 150bp, was carried out on the Illumina Hiseq X-ten system. Each experimental group consisted of two biological replicates.

### Ribo-seq data analysis

Ribo-seq data analysis was conducted as previously reported^44,73^. In brief, Seqprep was used to remove adapters and low-quality reads. Trimmomatic was used to further remove low quality reads. Reads were first aligned to rRNA and tRNA reference (hg38) followed by aligning to the genome (hg38) using bowtie2^75^. FeatureCounts was used to generate count files and Limma was used for differential analysis. Ribotoolkit^76^ was used to analyze codon occupancy and pausing. Translational efficiency was calculated by normalizing the Ribo-seq reads to the RNA-seq reads followed by differential expression analysis with Limma-voom using Riborex (https://github.com/smithlabcode/riborex). Codon analysis was conducted as previously reported^44,45,77^.

### Mass spectrometry detection of RNA modifications

Liquid chromatography tandem mass spectrometry (LC-MS/MS) was conducted as previously reported with modifications^44^. Here, we used an ultrafast high throughput analysis method that takes 10 minutes per sample to analyze > 45 mammalian modifications. In brief, 1.5∼3 μg tRNA enriched small RNA fraction (< 200 nt) was digested in a digestion buffer containing MgCl2 2.5 mM, Tris (pH 8) 5 mM, Coformycin 0.1 μg/ml, deferoxamine 0.1 mM, Butylated hydroxytoluene 0.1 mM, Benzonase 0.25 U/μL, Calf intestinal alkaline phosphatase (CIAP) 0.1 U/μL, and Phosphodiesterase I (PDE I) 0.003 U/μL for 6 hours at 37°C. Samples (600∼1200 ng total injected RNA) and standards were injected into Waters BEH C18 column (50 × 2.1 mm, 1.7 µm) coupled to an Agilent 1290 HPLC system and an Agilent 6495 triple-quad mass spectrometer. The LC system was conducted at 25 °C and a flow rate of 0.35 mL/min. Buffer A was composed of 0.02% formic acid (FA) in DDW. Buffer B was composed of 0.02% FA in 70% Acetonitrile. Buffer gradient was: 2 minutes 0% B, 4 minutes 5.7% B, 5.9 minutes 72% B, 6 minutes 100% B, 6.7 minutes 100% B, 6.75 minutes 0% B, and 10 minutes 0% B. List of transitions used are in (Supp. Table 2). Three biological replicates were analyzed per group.

### Reverse transcription-polymerase chain reaction (RT-PCR)

RNA was extracted using the miRNeasy mini kit per the manufacturer’s protocol. To proceed with cDNA synthesis, 1 µg of RNA was reverse transcribed using the High-Capacity RNA-to-cDNA™ kit (Thermo Fisher Scientific, Cat# 4387406) following the protocol instructions. Next, for the preparation of the reaction mix, each PCR reaction contained 1 µl of cDNA, 8 µl of Nuclease-free water, 10 µl of GoTaq® qPCR Master Mix (Promega, Cat# A6002), and 0.5 µl of each respective forward and reverse primers. Later, RT-PCR was performed in triplicate for the ANG gene (F: AAGATTCTTCCTCCTGGGAGCC), (R: CCCGTCTCCTCATGATGCTTT) as follows: 1 minute at 95°C then forty rounds at 95°C for 15 seconds, and 60°C for 1 minute. Finally, the fold change in expression was determined by calculating 2^(−ΔΔ CT)^78^, with beta-Actin as a reference gene (F: CATGTACGTTGCTATCCAGGC), (R: CTCCTTAATGTCACGCACGAT).

### Seahorse assay

We conducted seahorse assay using the Seahorse XF96 analyzer (Agilent, CA, USA) and Mito-stress test kit (Agilent, Cat# 103015-100), along with Seahorse XFe96/XF Pro FluxPak (Agilent, Cat# 103792-100), and Seahorse XF DMEM medium (without Phenol Red/pH 7.4/with HEPES/500 mL, Agilent, Cat# 103575-100) to quantify oxygen consumption rate (OCR) and extracellular acidification rate (ECAR). Initially, 35,000 cells/well were seeded in 96-well cell culture plates on day one, while the XFe96 sensor cartridge was hydrated in a 200 µL calibration medium. On the following day, the cell medium was replaced with DMEM, and cells were incubated at 37℃ in a CO₂-free incubator for 1 hour. During this period, 1 µM Oligomycin, carbonyl cyanide-4 (trifluoromethoxy) phenylhydrazone (FCCP) at 0.5 µM, Antimycin A, and Rotenone were loaded into the drug ports of the cartridge. Following loading, the sensor plate was placed in the analyzer for calibration. After calibration, the cell culture plate was loaded, and the analysis was initiated with the following program:

Set: Mixture 2 minutes, Measure 3 minutes, 4 sets in all steps. Ports: (A: Oligomycin / B: FCCP / C: Antimycin/Rotenone).

### Western blotting

Western blotting was carried out as previously mentioned^6^. In summary, cells were homogenized in T-PER™ tissue protein extraction reagent (Thermo Fisher Scientific, Cat# 78510) containing Triton(R) X-100 (Nacalai Tesque, Cat# 35501-15) and cOmplete™ protease inhibitor cocktail (Roche, Cat# 4693116001). After centrifugation, the proteins were isolated from the supernatant and measured using the Brachidonic-acid assay kit (BCA) (Thermo Fisher Scientific, Cat# 23227). Next, on Mini-PROTEAN TGX gel (Bio-Rad, Cat# 4561096), equal proteins loads were separated and then transferred to Polyvinylidene difluoride membrane (PVDF) (Bio-Rad, Cat# 1704156). Afterward, membranes were then blocked in 5% Skim milk powder (Fujifilm, Cat# 190-12865) mixed with Phosphate buffered saline with tween® (PBS-T) (Takara, Cat# T9183). The chosen primary antibody was then incubated overnight at 4°C, followed by incubation at room temperature with the secondary antibody (IgG detector solution v2). Furthermore, using Pierce ECL substrate reagent (Thermo Fisher Scientific, Cat# 32106), the signal was detected on a ChemiDoc machine (Bio-Rad). Finally, to normalize the protein expression, the Anti-beta actin antibody was reprobed after stripping the membranes using Restore™ stripping buffer (Thermo Fisher Scientific, Cat# 21059).

### List of antibodies used

- Angiogenin antibody (1:1000 dilution, Thermo Fisher Scientific, Cat# PA5-118948).
- Beta actin antibody (1:2000 dilution, Cell signaling, Cat# 4970S).
- EIF2a antibody (1:1000 dilution, Cell signaling, Cat# 9722).
- FLAG-M2 antibody (1:1000 dilution, Sigma Aldrich, Cat# F3165).
- G3BP1 antibody (1:1000 dilution, Invitrogen, Cat# PA5-29455).
- IgG detector solution V2 (1:2000 dilution, Takara, Cat# T7122A-1).
- Mouse IgG (1:1000 dilution, Cell signaling, Cat# 7076S).
- Puromycin antibody (1:2000 dilution, Millipore, Cat# MABE343).
- P38 antibody (1:1000 dilution, Cell signaling, Cat# 9212).
- p-P38 antibody (1:1000 dilution, Cell signaling, Cat# 4511s).
- p-EIF2a antibody (1:1000 dilution, Cell signaling, Cat# 3398).
- P70S6K antibody (1:1000 dilution, Cell signaling, Cat# 9202).
- p-P70S6K antibody (1:1000 dilution, Cell signaling, Cat# 9234).
- QTRT1 antibody (1:1000 dilution, GeneTex, Cat# GTX118778).
- QTRT2 antibody (1:1000 dilution, GeneTex, Cat# GTX123016).
- Total OXPHOS Human antibody cocktail (1:1000 dilution, abcam, Cat# ab110411).

### Puromycin incorporation assay

After being incubated in a complete growth medium with 10 μg/ml puromycin (Sigma Aldrich, Cat# P8833) for 30 minutes, the cells were harvested as previously described and subjected to western blotting analysis using puromycin antibody.

### Immunofluorescence assay and Confocal microscopy

Using Nunc™ Lab-Tek™ II Chamber Slide™ (Thermo Fisher Scientific, Cat# 154534PK) coated with collagen (Koken, Cat# IPC-50), 293T cells were fixed with 4% paraformaldehyde (Fujifilm, Cat# 162-16065) in Dulbecco’s phosphate-buffered saline (D-PBS) (Nacalai Tesque, Cat# 14249-24) for 10 minutes after the exposure to various stressors. D-PBS was used to wash the cells before and after each step. Then, cells were permeabilized using 0.1% Triton X-100 for 20 minutes at RT and blocked with 2% Bovine serum albumin (BSA) (Fujifilm, Cat# 010-25783) in D-PBS for 1 hour at RT. Next, G3BP1 or EDC4 (Santa Cruz Biotechnology, Cat# sc-374211) antibodies were used with 1% BSA and 0.1% Triton X-100 in D-PBS for overnight incubation at 4°C followed by a secondary incubation of Alexa Fluor 568 antibody (Invitrogen, Cat#A11036) or Alexa Fluor 488 antibody (Invitrogen, Cat# A11029) for 2 hours at RT. Afterward, cell nuclei were stained with 4’,6-Diamidino-2-Phenylindole, Dihydrochloride (DAPI) (1:500 dilution, Thermo Fisher Scientific, Cat# D1306) for 2 minutes at RT. Finally, cells were mounted with ProLong™ Diamond Antifade Mountant (Thermo Fisher Scientific, Cat# P36961). Live cell imaging was conducted with the MitoTracker™ Green FM (Invitrogen, Cat# M7514) and MitoTracker™ Deep Red FM (Invitrogen, Cat# M22426) using a glass bottom dish (Matsunami, Cat# D11131H). Moreover, confocal microscopy with Fluoroview v6 software (Olympus FV3000) was utilized to obtain images.

### Data visualization and statistical analysis

Heatmaps, clustering and Pearson’s correlation analysis were conducted using Morpheus (https://software.broadinstitute.org/morpheus). For the analysis of western blotting, the ImageJ software^79^ was used where data were presented as means from three independent experiments. Comparisons were performed with unpaired Student t-test (two-tailed) or Two Way ANOVA using the Prism Graph version 9.3 software. The statistical significance was set at P < 0.05 and fold change of more than 1.5 folds. Results are denoted by asterisks in figures (*: significant when compared to the control / **: significant when compared to Mock or another group).

## Supporting information

Suppmenetary table 1

Suppmenetary table 2

## Acknowledgment

The authors report no conflict of interest nor any ethical adherences regarding this work.

## Author contributions

S.A.: Study design. Performed all experiments. Data analysis and interpretation. Manuscript writing. K.T.: Seahorse analysis. A.M.: Critically revised the manuscript. T.J.B.: Critically revised the manuscript. Codon analysis algorithm development. P.C.D.: Critically revised manuscript. S.R.: Study conception and design. Funding acquisition. Supervision. Mass spectrometry analysis. Bioinformatics analysis. Data analysis and interpretation. Manuscript writing. Administration. K.N.: Critically revised the manuscript. Supervision. Administration.

## Data availability

Sequencing data presented herein are deposited in the sequence read archive database with accession numbers: PRJNA1002608 (small RNA-seq) and PRJNA1061219 (RNA-seq and Ribo-seq).

