## Supplemental data for "Angiogenin regulates mitochondrial stress and function via tRNA-derived fragments generation and impacting tRNA modifications"

### Supplementary data

#### Supplementary tables information:

**Supplementary table 1:** The transition list of LC-MS/MS analysis of tRNA modifications.

**Supplementary table 2:** Normalized peak area of tRNA modifications.

#### Supplementary figures [1-12]

**Supplementary figure 1: A:** Quantification of puromycin incorporation assay after stress. Asterisk: fold change > 1.5, $p < 0.05$. **B-C:** Puromycin incorporation assay time course of arsenite. **D-F:** Phosphorylation rate percentage of ISR and RSR markers. **G:** Quantification of SYBR gold staining. Asterisk: fold change > 1.5, $p < 0.05$.

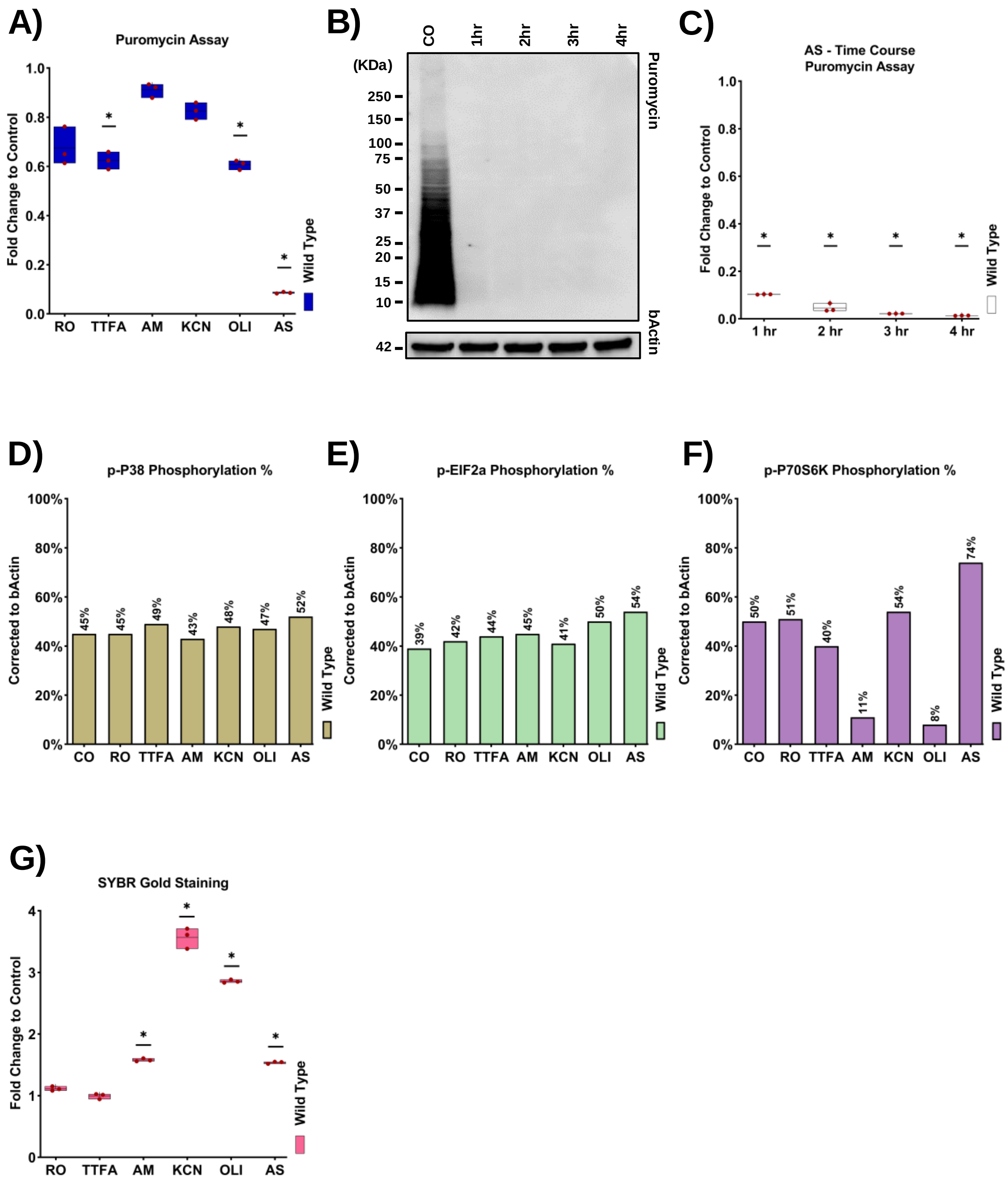

**Supplementary figure 2: A:** Sanger sequencing for the successfully inserted oligos. **B:** Puromycin incorporation assay after stress. Asterisk: fold change > 1.5, p < 0.05. **C:** Quantification of puromycin incorporation assay. **D-L:** Quantification of ISR and RSR markers.

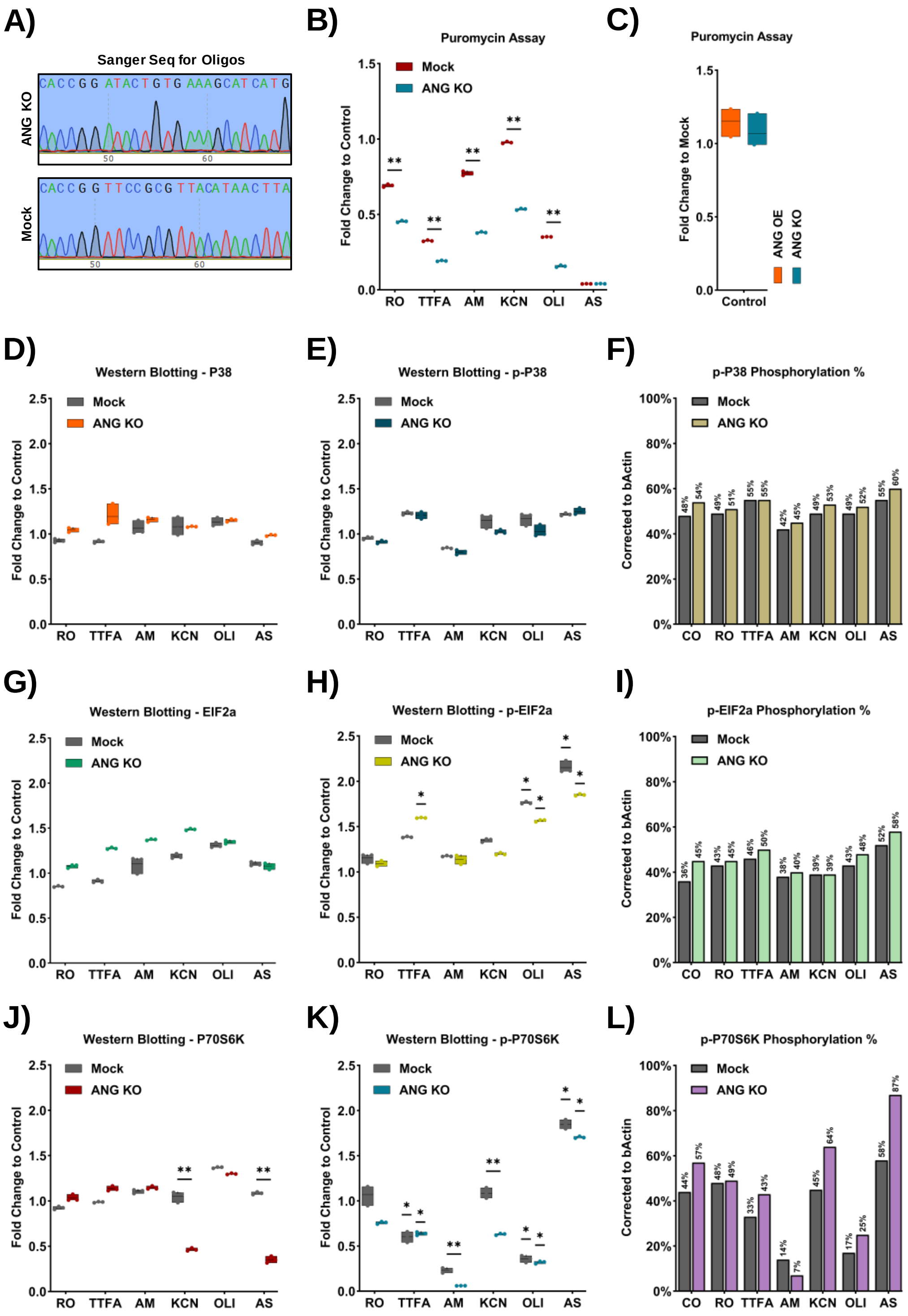

**Supplementary figure 3: A:** Quantification of SYBR gold staining. Asterisk: fold change > 1.5, p < 0.05. **B-I:** Quantification of seahorse assay. Asterisk: fold change > 1.5, p < 0.05. **J:** Quantification of OXPHOS-related proteins in all mitochondrial complexes. Asterisk: fold change > 1.5, p < 0.05. **K:** Immunofluorescence of Mitotracker green and deep red, 60X magnification, scale bar 20 µm. **L:** Immunofluorescence of EDC4 (green), counterstained with DAPI (blue), 60X magnification, scale bar 20 µm.

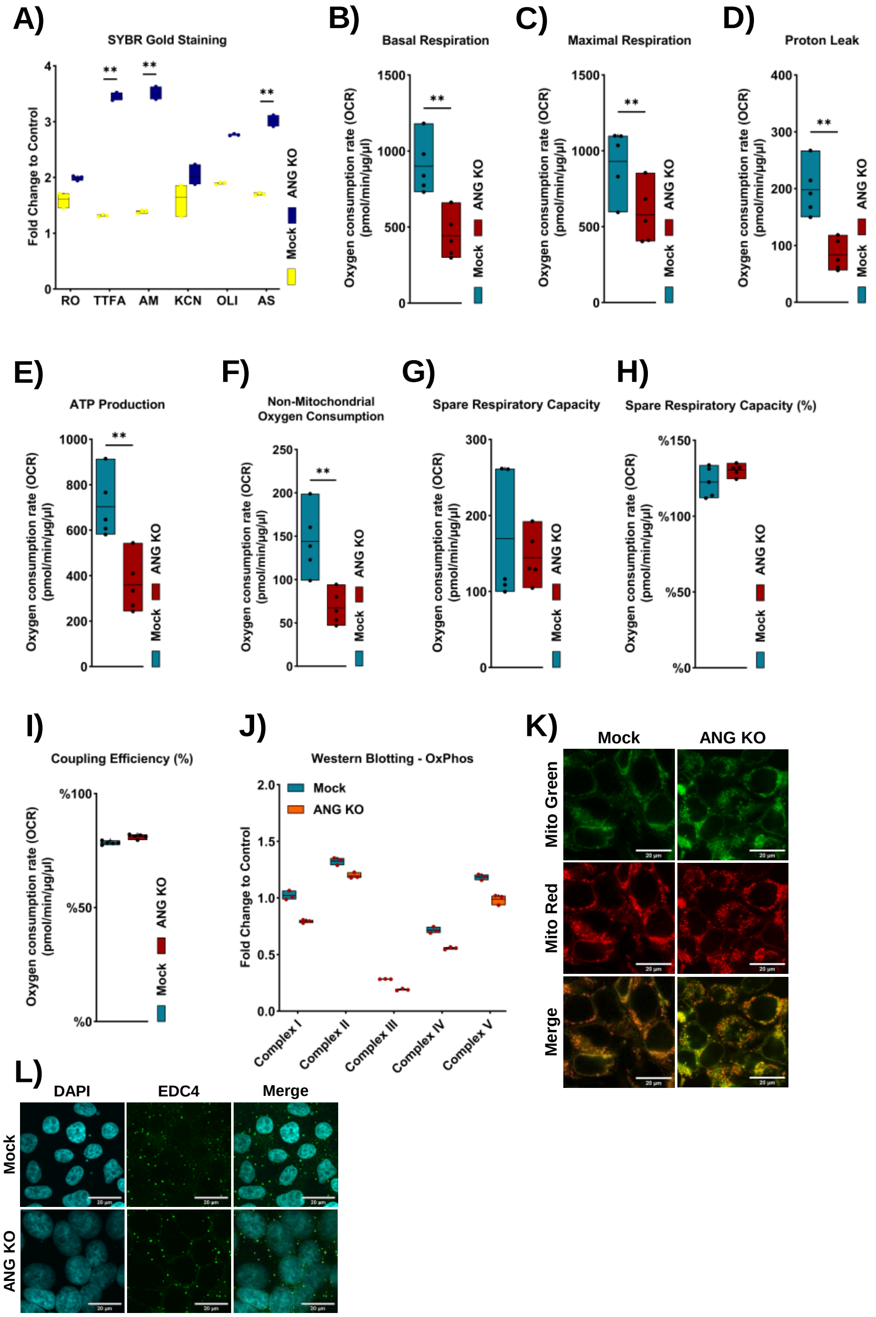

**Supplementary figure 4: A-D:** Volcano plots of the Small RNA sequencing dataset for ANG-OE.

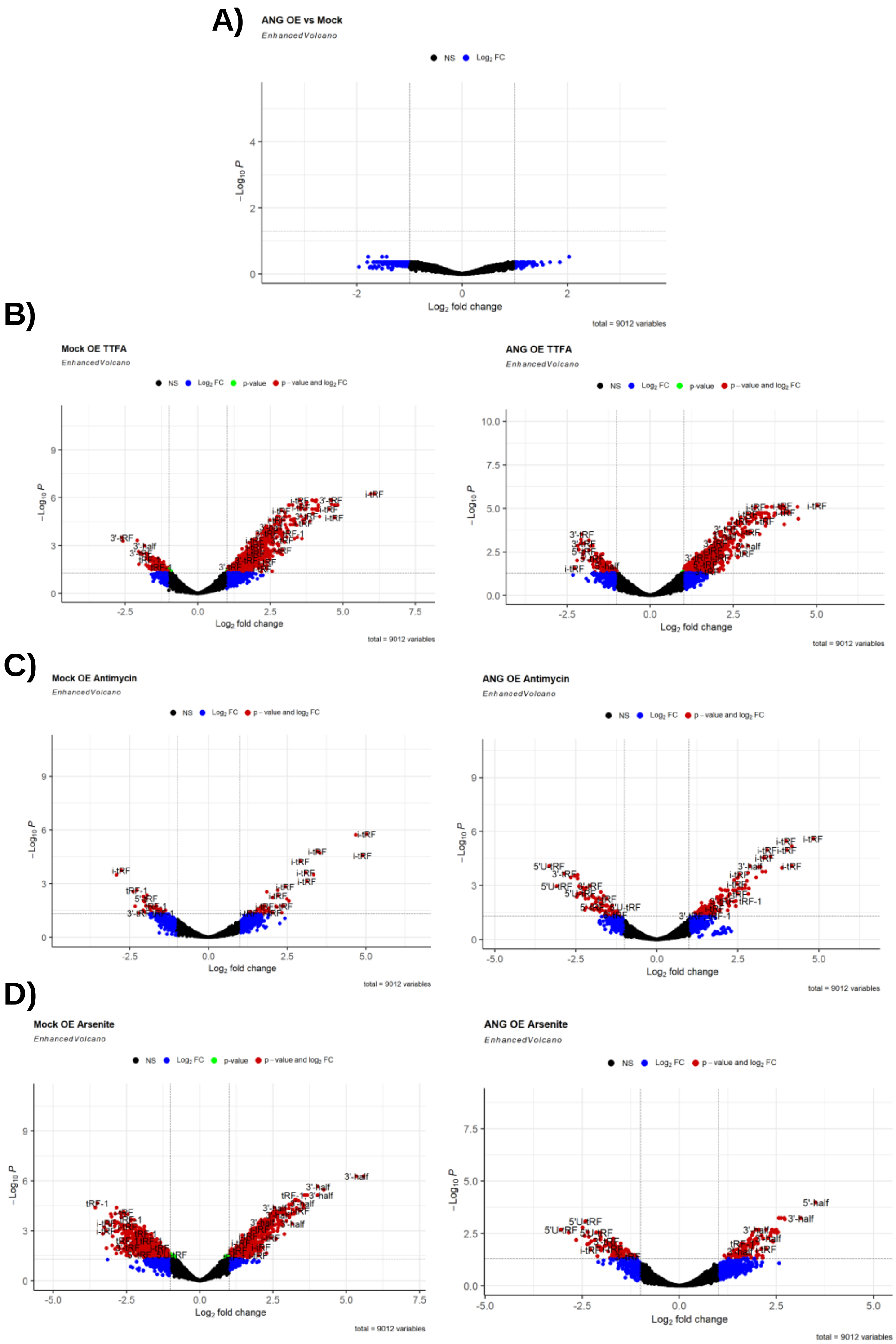

**Supplementary figure 5: A-D:** Volcano plots of the Small RNA sequencing dataset for normalized ANG- (OE vs KO).

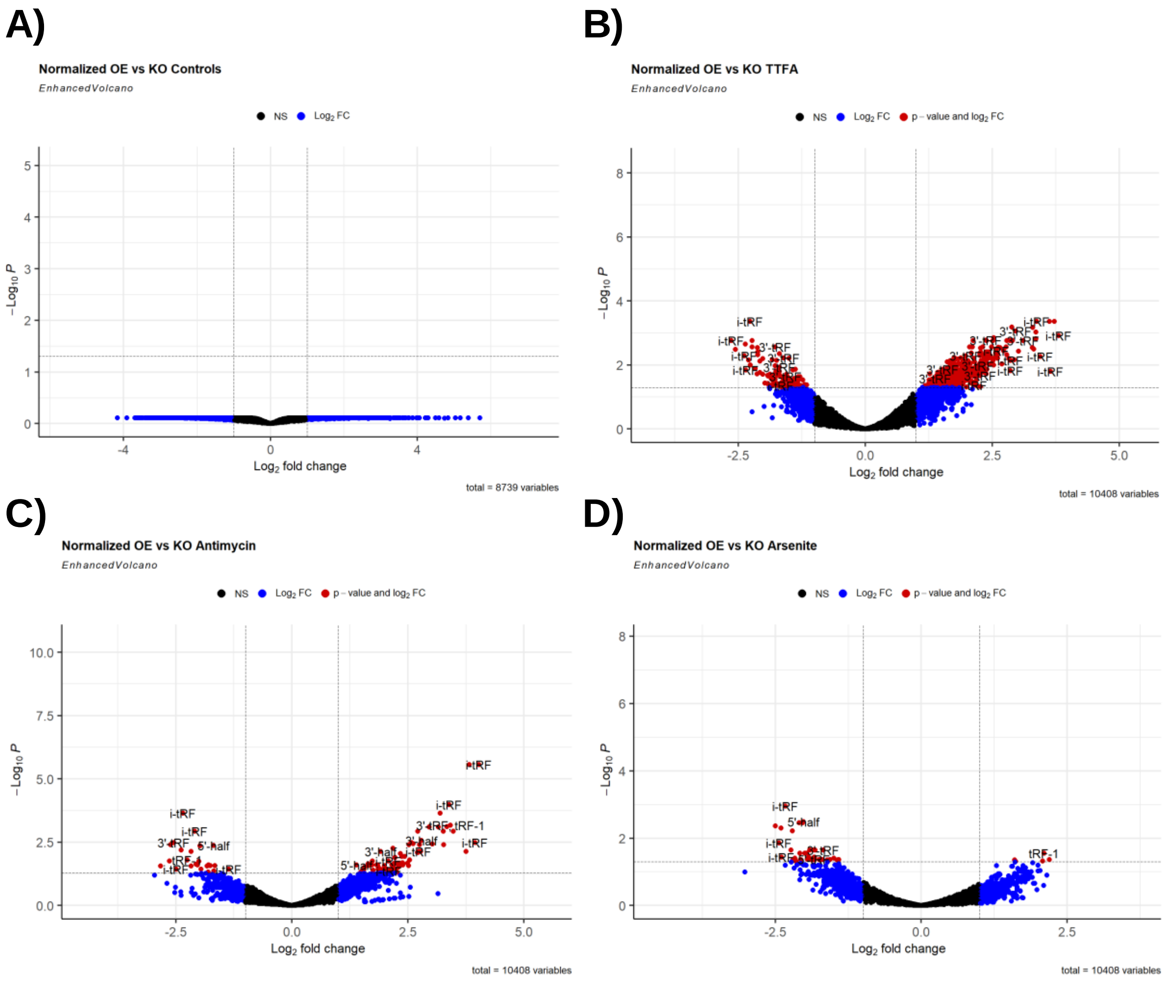

**Supplementary figure 6: A:** Motif analysis of significant tsRNAs after stress. **B:** Motif analysis comparison between AM and other stresses.

**A)**

| Stress | Motif | Regular expression | Logo | E-value |
|---|---|---|---|---|
| TTFA | TSMGAGAA | T[GC][CA]GAGAA | 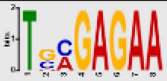 | 4.8e-044 |
| | GGTKBTAR | GGT[TG][GCT][TG]A[GA] | 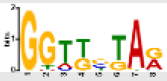 | 1.9e-038 |
| | TTGGTCGT | TTGGTCGT | 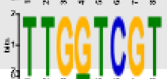 | 6.1e-017 |
| AM | GRGTTCGA | G[AG]GTTCGA | 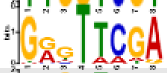 | 3.2e-039 |
| | RAGGTYGY | [GA][AG]G[GA]T[TC]G[TC] | 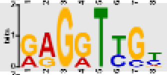 | 1.7e-005 |
| | ATTGGVCT | AT[TG][GA]G[ACG]CT | 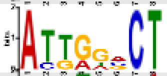 | 5.9e-003 |
| AS | GGGTTCGA | G[GA]GTTCGA | 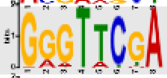 | 1.0e-143 |
| | AATCCATT | AATCCA[TA]T | 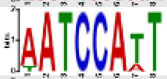 | 1.0e-026 |
| | TCCYCGTC | TCC[TC][CG]GT[CT] | 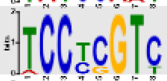 | 9.3e-016 |
| | VTAGTGGT | [AGC]TAGTGGT | 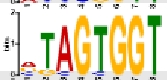 | 2.8e-014 |
| | TAATCTGA | TAATCTGA | 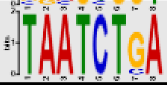 | 1.1e-010 |

**B)**

| Comparison | Motif | Regular expression | Logo | E-value |
|---|---|---|---|---|
| AM vs TTFA | RGTTCGAT | [AG]G[TG]T[CT][GA]A[TAG] | 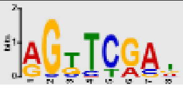 | 1.6e-025 |
| | RGRTTGTG | [GA][GC][AG]TT[GA]TG | 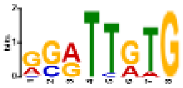 | 1.1e-012 |
| AM vs AS | TTCGADYC | TTCGA[GAT][CT]C | 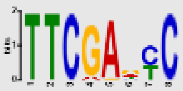 | 4.6e-033 |
| | ADCRACCT | [AC][GAT]C[AG]ACCT | 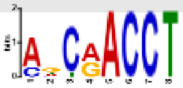 | 2.0e-007 |
| | CCCACAAY | [CG][CT]CA[CT]AA[TC] | 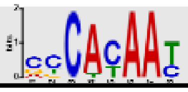 | 2.7e-010 |

**Supplementary figure 7: A-B:** Motif analysis for RNA binding proteins using the top motifs in the previous comparison analysis with AM.

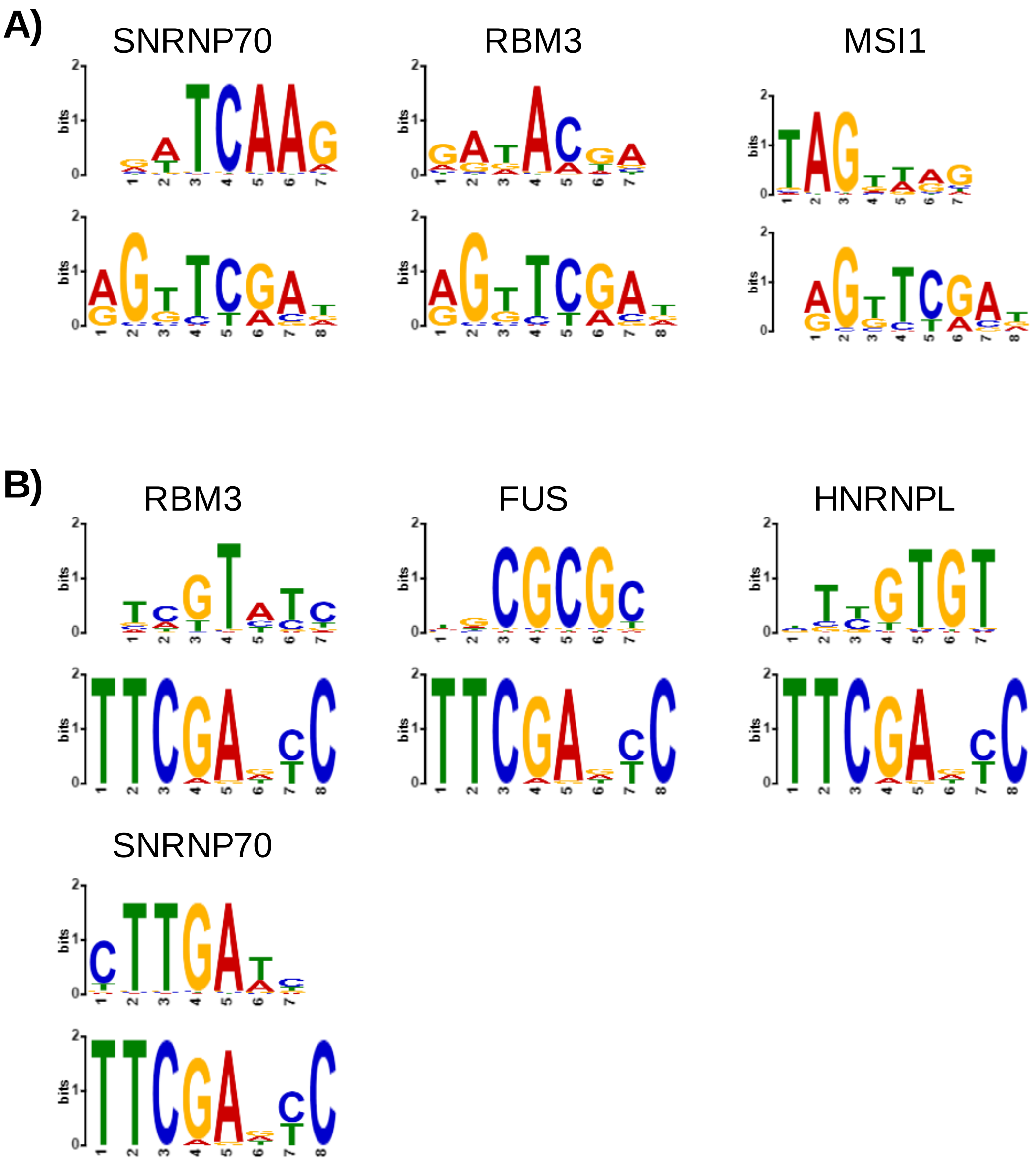

**Supplementary figure 8: A:** Analysis of tRNA modifying enzymes expression related to tRNA-Q and yW modifications in RNA-seq, translation efficiency, and Ribo-seq datasets. **B-C:** Western blotting of QTRT1 and QTRT2 in the ANG-KO cells. Asterisk: fold change $> 1.5$, $p < 0.05$. **D:** Volcano plot of mature tRNAs expression in ANG-KO vs Mock-KO cells.

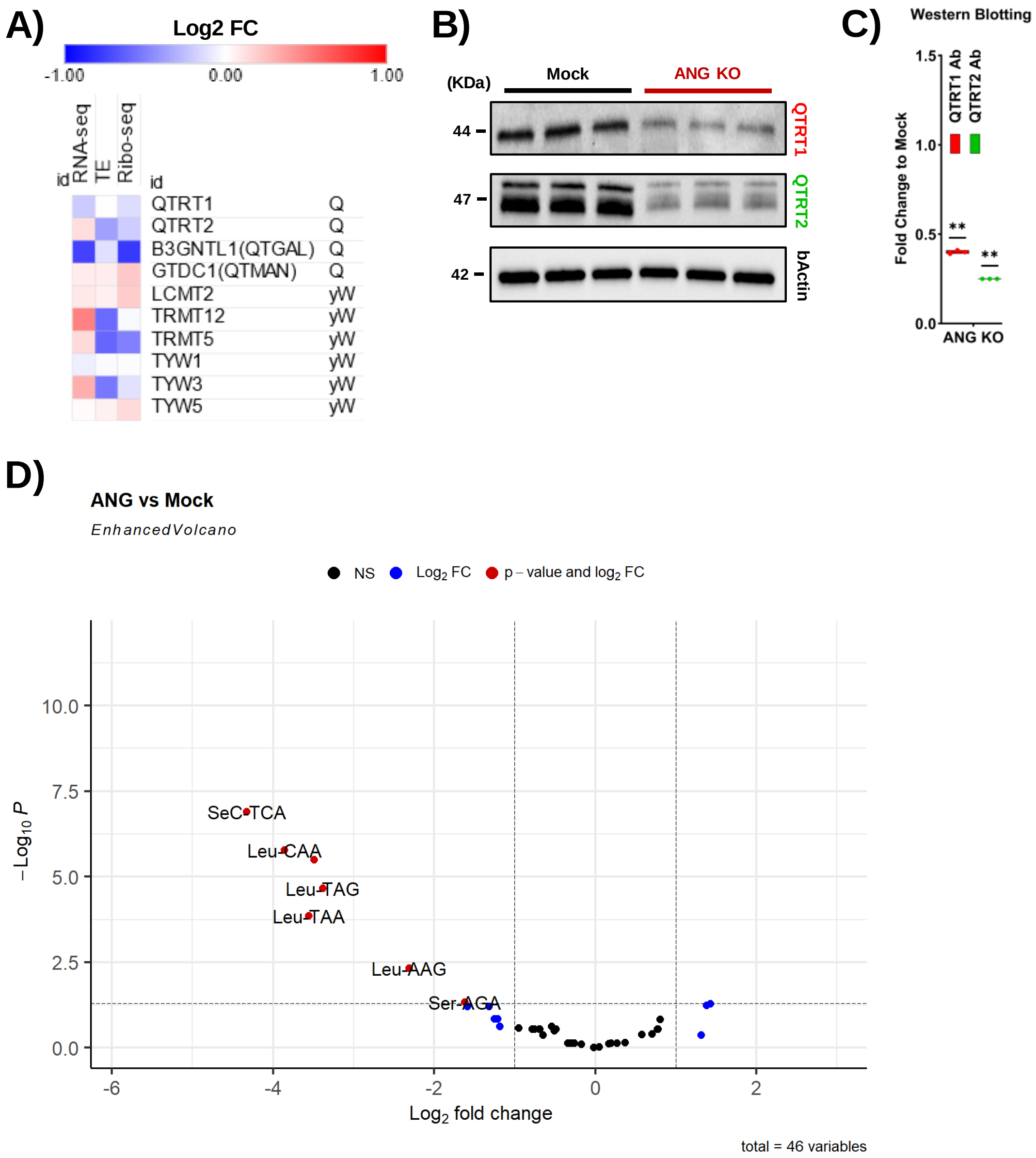

**Supplementary figure 9: A:** Heatmaps of tRNA modifications of wild-type cells. **B:** Heatmaps of tRNA modifications of Mock-OE. **C:** Heatmaps of tRNA modifications of ANG-OE. **D-K:** Area ratio of significant tRNA modifications. Asterisk: fold change > 1.5, p < 0.05.

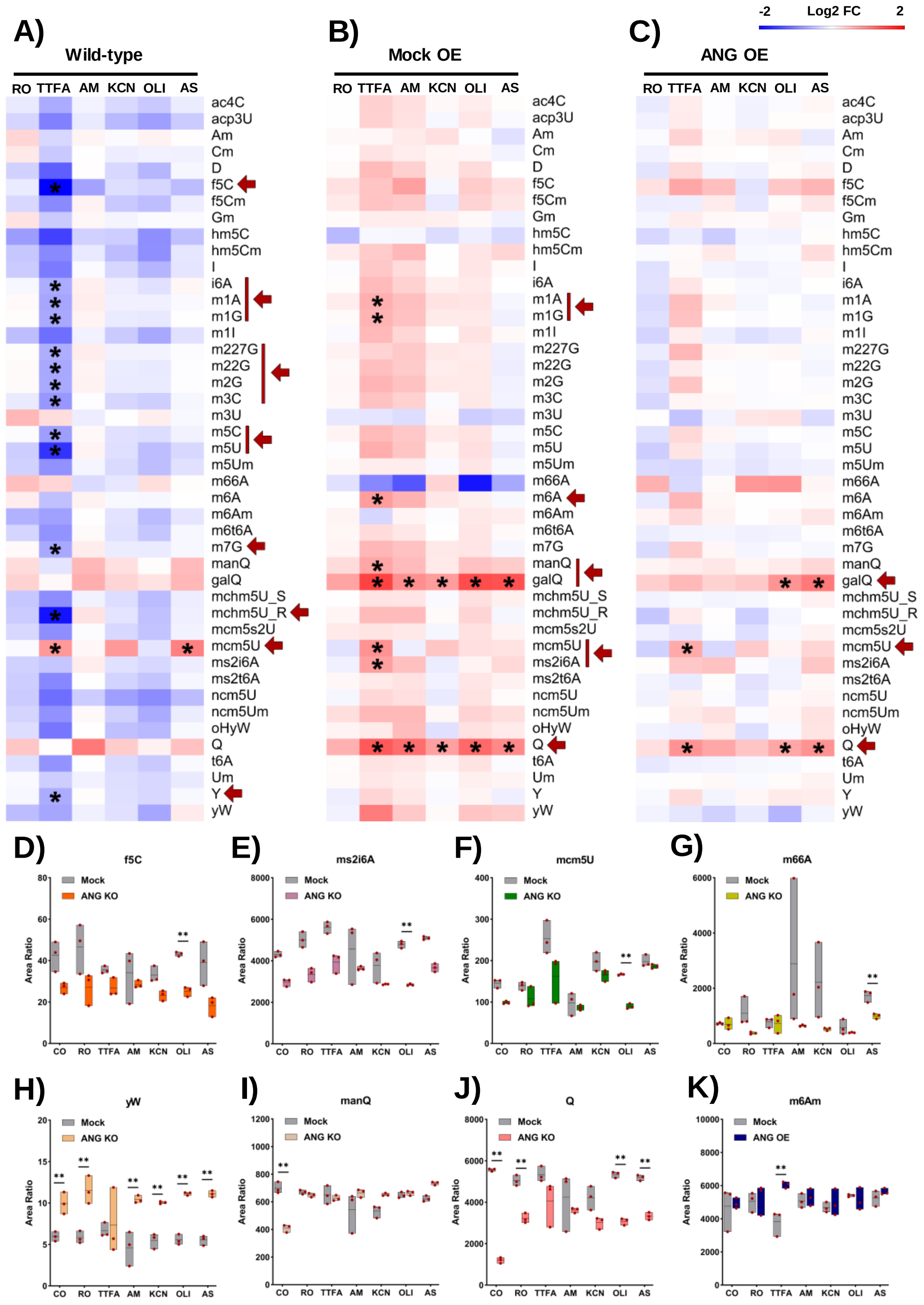

**Supplementary figure 10: A:** RA and Kegg of the RNA sequencing dataset. **B:** RA and Kegg of the Ribosome profiling dataset. **C:** RA and Kegg of the Translation efficiency dataset.

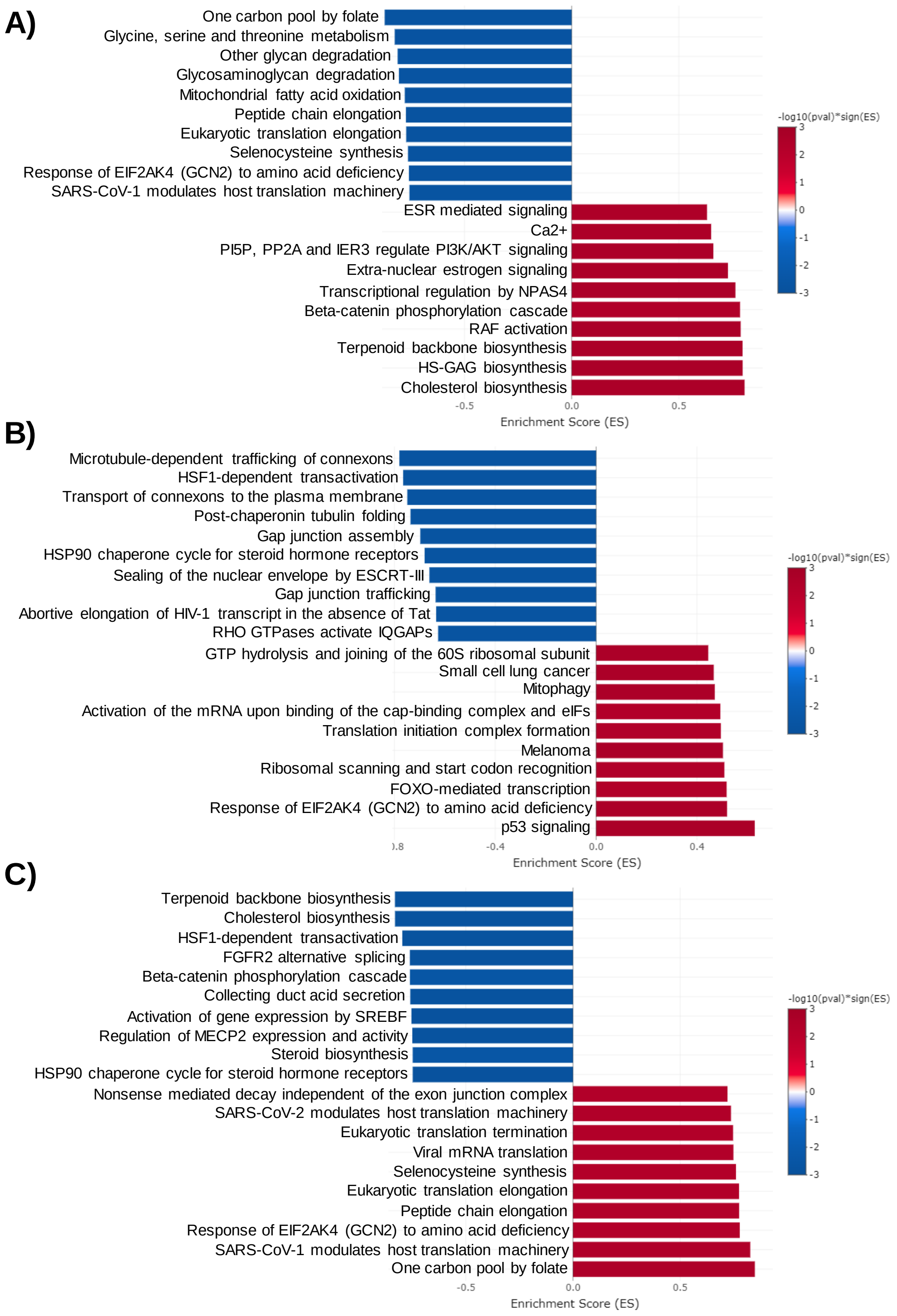

**Supplementary figure 11: A:** Isoacceptors codon frequency analysis of the Translation efficiency dataset. **B:** Total codon count analysis of the Translation efficiency dataset.

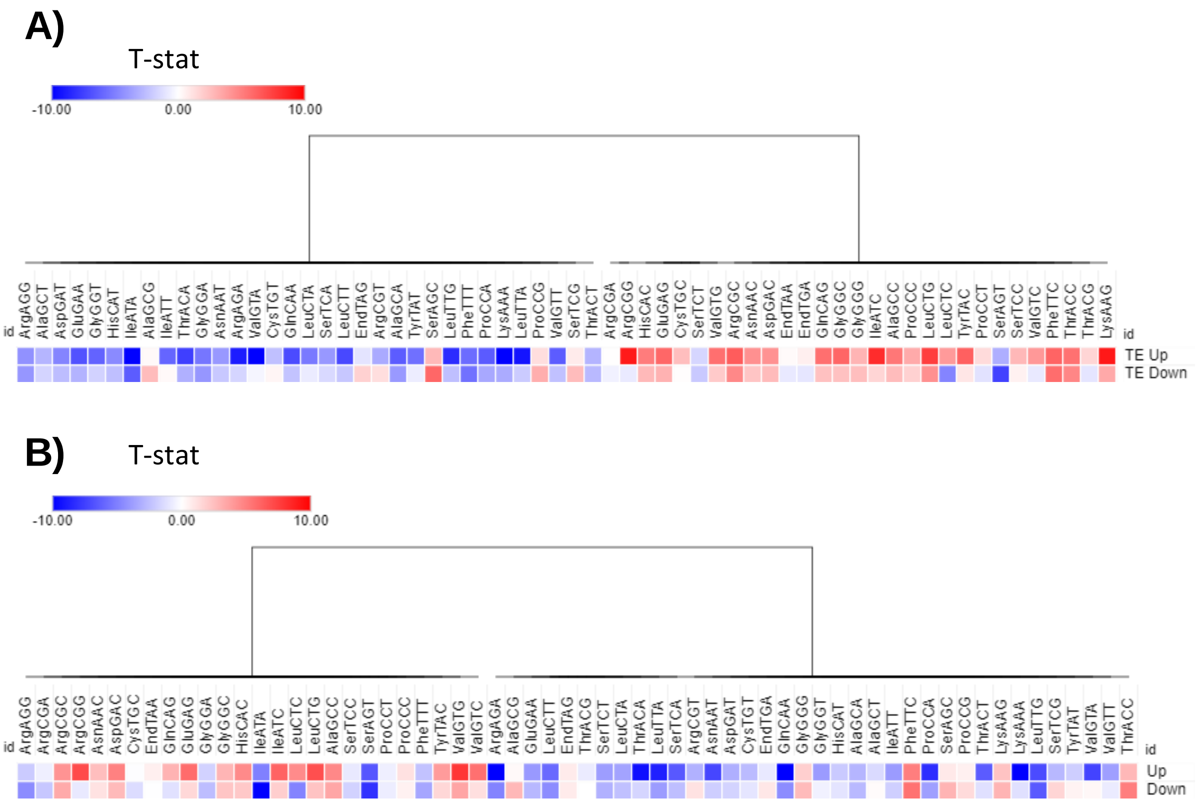

**Supplementary figure 12: A:** Sanger sequencing for the successfully inserted shRNAs. **B:** Colony PCR for the successfully inserted shRNAs.

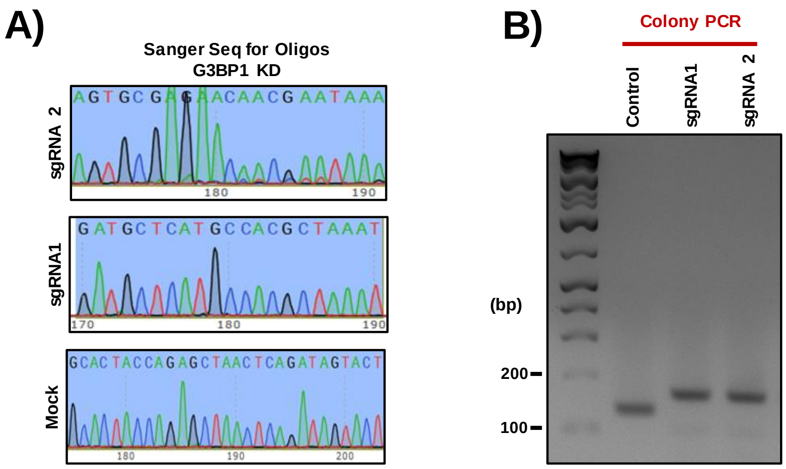
